# A conserved subset of cold tumors responsive to immune checkpoint blockade

**DOI:** 10.1101/2024.03.06.583752

**Authors:** Jade Moore, Jim Gkantalis, Ines Guix, William Chou, Kobe Yuen, Ann A. Lazar, Mathew Spitzer, Alexis J. Combes, Mary Helen Barcellos-Hoff

## Abstract

**Background:** The efficacy of immune checkpoint blockade (ICB) depends on restoring immune recognition of cancer cells that have evaded immune surveillance. At the time of diagnosis, patients with lymphocyte-infiltrated cancers are the most responsive to ICB, yet a considerable fraction of patients have immune-poor tumors.

**Methods:** We analyzed transcriptomic data from IMvigor210, TCGA, and TISMO datasets to evaluate the predictive value of βAlt, a score representing the negative correlation of signatures consisting of transforming growth factor beta (TGFβ) targets and genes involved in error-prone DNA repair. The immune context of βAlt was assessed by evaluating tumor-educated immune signatures. An ICB-resistant, high βAlt preclinical tumor model was treated with a TGFβ inhibitor, radiation, and/or ICB and assessed for immune composition and tumor control.

**Results:** Here, we show that high βAlt is associated with an immune-poor context yet is predictive of ICB response in both humans and mice. A high βAlt cancer in which TGFβ signaling is compromised generates a TGFβ rich, immunosuppressive tumor microenvironment. Accordingly, preclinical modeling showed that TGFβ inhibition followed by radiotherapy could convert an immune-poor, ICB-resistant tumor to an immune-rich, ICB-responsive tumor. Mechanistically, TGFβ blockade in irradiated tumors activated natural killer cells that were required to recruit lymphocytes to respond to ICB. In support of this, natural killer cell activation signatures were also increased in immune-poor mouse and human tumors that responded to ICB.

**Conclusions:** These studies suggest that loss of TGFβ competency identifies a subset of cold tumors that are candidates for ICB. Our mechanistic studies show that inhibiting TGFβ activity converts high βAlt, cold tumors into ICB-responsive tumors via NK cells. Thus, a biomarker consisting of combined TGFβ, DNA repair, and immune context signatures provides a means to prospectively identify patients whose cancers may be converted from ‘cold’ to ‘hot,’ which could be exploited for therapeutic treatment.

- **What is already known on this topic** – *For some cancer patients, response to ICB provides durable tumor control. Current biomarkers are insufficient to reliably predict the immunotherapy response for most patients, particularly those whose tumors lack lymphocytic infiltration*.
- **What this study adds** – *The βAlt score, which reports a DNA damage deficiency caused by lack of TGFβ signaling, predicts response to ICB in clinical trial data from IMvigor210 metastatic bladder cancer patients and for melanoma patients. Notably, transcriptomic assessment of the immune context shows that these are immune-poor, so-called “cold” tumors. Preclinical modeling indicates that alleviating TGFβ inhibition of NK cells is critical to relieving immunosuppression*.
- **How this study might affect research, practice or policy** – *Our work identifies a novel tumor phenotype consisting of cancer cells that have lost TGFβ signaling and gained error-prone DNA repair embedded in a TGFβ rich, immune-poor microenvironment, which is conserved across cancer types in humans and among preclinical tumor models. Patients whose immune-poor tumors have high βAlt scores are strong candidates for ICB and radiotherapy combinations that may be further augmented by TGFβ inhibition. Hence, the βAlt score can be used to stratify immune-poor cancer patients for optimal therapeutic strategies.*

## Background

Understanding the specific mechanisms by which cancer has subjugated the immune system is essential for cancer immunotherapy to restore immune recognition. The canonical example is immune checkpoint blockade (ICB) targeting programmed cell death protein one or programmed death-ligand 1 (PD-1/PD-L1) or cytotoxic T-associated lymphocyte four, which are now the standard of care for many adult cancers, yielding significant and durable clinical responses in a subset of patients ^1^. Patient selection for ICB is primarily guided by histopathological assessment of PD-L1 expression and/or the spatial distribution or density of tumor-infiltrating lymphocytes ^2^. High tumor mutation burden is also associated with response to ICB in several settings ^3, 4^ and is an FDA-approved biomarker for ICB in certain cancers ^5, 6^. Experimentally, cancers are frequently characterized in terms of immune composition, as immune-rich, ‘hot’ or infiltrated versus immune deserts, ‘cold,’ or devoid of lymphocytes ^7–10^. Various methods, including digital pathology, transcriptional signatures, and immunohistochemistry, report lymphocytic infiltration ^11, 12^. Patients with tumors infiltrated with lymphocytes are considered good candidates for ICB response, whereas those whose tumors lack lymphocytes are not.

A notable success for predicting patient response stemmed from recognition that patients with mismatch repair deficient (dMMR) tumors were enriched in some early ICB trials ^13^. This led to the NICHE-2 trial in which most colorectal cancer patients with dMMR responded to neoadjuvant ICB therapy; moreover, the association of dMMR with response rate and duration to ICB is also predictive in 12 other tumor types ^14^. Misrepair of intrinsic or extrinsic DNA damage in the dMMR context is believed to provoke immune recognition through increased neoantigen production, release, and presentation ^15, 16^. Consistent with this, dMMR tumors are highly inflamed due to cytosolic DNA sensing mediated by cGAS/STING that elicits type I interferon (IFN) signaling to provoke lymphocyte recruitment ^17^.

Transforming growth factor-beta (TGFβ) is a high value target in cancer therapy because it promotes a permissive tumor microenvironment (TME) and invasive cancer phenotypes while suppressing anti-tumor immunity ^18, 19^. Notably, TGFβ also has a crucial role in regulating the DNA damage response ^20^. TGFβ positively regulates canonical DNA repair by homologous recombination and non-homologous end-joining and suppresses repair by error-prone alternative end-joining (alt-EJ) ^20–22^. This relationship is reported by gene signatures consisting of TGFβ signaling targets and genes necessary for alt-EJ DNA repair, which are significantly anti-correlated across most tumor types and represented by a score termed βAlt that predicts response to genotoxic therapy ^23, 24^. Cancers with a high βAlt score are those in which TGFβ signaling is compromised and express high levels of alt-EJ genes. Consistent with defective DNA repair, high βAlt tumors have a greater fraction of the genome altered, are characterized by a specific indel mutation, and are more responsive to chemoradiation ^23^.

Since genomic instability is a hallmark of cancer that can induce type I interferon to activate innate immunity ^25^, it has been proposed that specific DNA repair deficiencies associated with response to genotoxic therapies may also predict response to immunotherapy ^26^. Given that TGFβ determines DNA repair competency, we hypothesized that low βAlt tumors, exhibiting high expression of TGFβ target genes, have obligate TGFβ activity and would be immune-poor. In contrast, high βAlt cancers with impaired TGFβ signaling resulting in poor DNA repair, like those with dMMR, would be infiltrated. We tested this hypothesis by concurrently determining βAlt and the immune context for tumors of patients who responded to immunotherapy in clinical trials. High βAlt strongly correlated with ICB response in both bladder cancer and melanoma patients but was unexpectedly associated with immune-poor cancer. Indeed, the association between high βAlt and immune-poor cancers was highly conserved across human cancer types and in mouse cancers. Mechanistic studies in preclinical models showed that TGFβ blockade in irradiated tumors activated natural killer cells that were required to recruit lymphocytes and respond to ICB. Natural killer cell activation was also evident in immune-poor human tumors that responded to ICB.

### These studies identified a previously unrecognized subset of immune-poor patients whose

βAlt status predicts ICB response.

## Methods

### Animals

All animal experiments were performed at the University of California, San Francisco (UCSF, San Francisco, CA, USA). The protocols for animal husbandry and experiments were conducted with approval from the UCSF institutional review board (AN142057-01A) and adhered to the National Institutes of Health Guide for the Care and Use of Laboratory Animals. Eight-week-old BABL/cJ *Mus musculus* (RRID: IMSF_JAX:000651) from Jackson Laboratory (Sacramento, CA, USA) were purchased and housed five per cage, fed Lab Diet #5001 Rodent Formular (Purina Animal Nutrition LLC), and supplied water *ad libitum*.

### Mammary tumor-derived transplants

For the treatment experiments, 8-10-week-old mice were transplanted orthotopically with F2 mTDT from family F previously described ^27^. For all experiments, mice were palpated three times a week until the tumor reached 2 × 2, and then daily upon randomization into treatment groups when the tumor reached 65-100mm^3^. Tumor burden was calculated based on 0.52 x w × l^2^ and used to randomize mice to treatment groups as follows: Sham IgG, anti-programmed death ligand (PD-L1), TGFβ inhibition with small molecule IPW or single-dose radiation treatment, and dual and triple combinations.

All agents were administered intraperitoneally 24 hours before irradiation at the indicated schedule for up to three weeks. Anti-PD-L1 (Roche, Inc.) was administered once a week (10 mg/kg, PD-L1 9708, 6E11), TGFβ small molecule inhibitor IPW-5371 (Innovation Pathways, Inc.; designated as IPW) was administered daily (20 mg/kg), or IgG control antibody (Bioxcell BP0083) 3 times per week (25 mg/kg). Tumors were irradiated with 10 Gy using a small animal radiation research platform (SARRP, XStrahl) and individualized plans (Muriplan, XStrahl) based on arc beam computerized tomography (CT). For in vivo depletion of NK cells, mice received 50 mg/ml i.p. of asialo-GM1 polyclonal antibody (eBioscience^TM^, RRID: AB_10718540) two days before and one day after irradiation.

Mice were monitored daily and sacrificed for tumor ulcers or weight loss greater than 15% of body weight in accordance with the UCSF Institutional Animal Care and Use Committee guidelines. Blinded technicians performed tumor measurements and collection for the treatment groups. Each mouse was assigned to one of two categories based on tumor growth during the first 7-days post-treatment: responders (R) and non-responders (NR). Mice whose tumor volumes did not double from the start of treatment until seven days post-treatment were classified as responders, whereas mice whose tumors doubled daily and failed to meet these criteria were classified as non-responders. Sham-irradiated, IgG-treated mice were excluded from therapeutic response estimation.

### RNA sequencing

For bulk RNA sequencing, fragments (∼50 mg) of liquid nitrogen-preserved mTDT were shipped on dry ice to Q^2^ Solutions | EA Genomics to isolate total RNA using proprietary methods. The quantity and quality of the specimens were assessed using an Agilent Bioanalyzer and NanoDrop analysis. Total RNA samples that met the quality control metrics were enriched for mRNA, fragmented, and converted into indexed cDNA libraries for Illumina sequencing. The generated cDNA libraries were quantified by qPCR using primers specific for Illumina sequencing adapters, and TruSeq stranded mRNA in a 30 M 50 bp PE assay. Raw sequencing data were obtained in fastq format. Read mapping was performed using the Rsubread package (RRID: SCR_016945) against *Mus musculus* GRCm38 and the human GRCh38 genome (<u>28</u>). Gene expression data archived in the Gene Expression Omnibus (GEO, RRID: SCR_005012) are pending.

### Multispectral flow cytometry

Tumor and blood cells treated under the conditions described were counted with 1:1 Trypan blue/cells and blocked with 0.25 mg TruStain FcX PLUS antibody (BioLegend Cat#156603, clone S17011E) per 10^6^ cells in 0.1mL cell staining buffer (CSB, 0.5% fetal bovine serum in PBS) for 30 minutes on ice. Cells were then washed with PBS and incubated with LIVE/DEAD Zombie Aqua (1:2000, BioLegend Cat#423101) in 0.02mL PBS and subsequently stained in 0.08 mL CSB using a panel of optimized fluorophore-conjugated cell surface antibodies (*Supplemental Table 1*) for 1 hour at 4°C in the dark. Cells were then washed 1X with 2 mL CSB, followed by centrifugation at 350 × *g* for 5 min at 4°C. The supernatant was aspirated, and the cell pellet was in 0.5 ml/tube FluorFix buffer (BioLegend Cat#422101) in the dark for 1h at room temperature for fixation. For intracellular staining, the nucleus was washed and permeabilized by resuspending in 0.5mL 1X intracellular staining permeabilization wash buffer (BioLegend Cat#421002) diluted in distilled water, centrifuged at 350 × *g* for 5 min at 4°C, and incubated with fluorophore-conjugated intracellular antibodies for 30 min in the dark at room temperature. Cells were then washed 2X with 2 mL intracellular staining perm wash buffer and centrifuged at 350 × *g* for 5 min at 4°C before resuspending the cells in CSB (0.5 mL) and analyzed on a 3-laser Cytek® Northern Lights spectral cytometer (Fremont, CA USA). The experimental samples were unmixed using appropriate cell-based single-color controls in SpectroFlo® (Cytek Biosciences). Data analysis was performed using the FCS Express 7 Research Edition De Novo software (Pasadena, CA, USA), employing suitable gating strategies (*Supplemental Table 2*) for each tissue. Each surface marker was expressed as median fluorescence intensity (MFI). An unsupervised analysis was performed using the dimensionality reduction algorithm t-distributed Stochastic Neighbor Embedding (t-SNE) with the Barnes-Hut approximation to identify and visualize populations. All FCS files were merged, and 100,000 live CD45+ cells were randomly downsampled from each sample to be subjected to the transformation algorithm to form clusters according to the expression levels of cell-surface phenotyping markers. Samples were grouped and overlaid according to treatment and response to visualize the differences in immune cell composition.

### Bioinformatics analyses

Gene expression heatmaps were constructed by hierarchical clustering using the R package ComplexHeatmap ^28^. Gene expression values were normalized z-scores for all heatmaps, Euclidian distance, and ward.D.2 clustering was used.

βAlt scores for preclinical tumors or patients were calculated based on the expression pattern using the previously published TGFβ and alt-EJ gene signatures ^23^, weighted as described ^24^. The composition of each signature is listed in *Supplemental Table 3*. In brief, the weight of each TGFβ gene and each alt-EJ gene was calculated by multiplying by its factor, and the βAlt weighted score was calculated as the sum of the weighted expression of the genes from each signature for each tumor. This βAlt score conveys the relative expression of each signature for each tumor.

Previously reported TeIS gene signatures ^29^ are listed in *Supplemental Table 4*. The feature gene signature scores for the 10 cell types included T cell, myeloid, CD90+CD44+ stroma, CD4, CD8, T regulatory cells, macrophages, monocytes, cDC1, and cDC2 ^29^.

### Datasets

The TCGA dataset was downloaded from the Genomic Data Commons portal from the EBPlusPlusAdjustPANCAN_IlluminaHiSeq_RNASeqV2.geneExp.tsv file. The downloaded gene expression values were trimmed as the mean of M values normalized, log2 transformed, and mean-centered per gene by converting them into z-scores. Primary solid tumor samples were analyzed. TCGA somatic mutations at the individual level were obtained following approval from the dbGaP Data Access Committee (Project #11689) ^30^.

The source code and processed data for IMvigor210 trial (NCT02108652) were accessed in IMvigor210CoreBiologies, a fully documented software, and package for the R statistical computing environment ^10^. The IMvigor010 dataset was from the European Genome-Phenome Archive under accession number EGAS00001004997 ^31^. The GSE78220 melanoma dataset (n=28 specimens) ^32^ and GSE91061 melanoma dataset (n=109 specimens with 58 on-treatment and 51 pre-treatment) from 65 patients ^33^ were downloaded from the Gene Expression Omnibus (GEO, RRID: SCR_005012) in January 2022 using the R package GEOquery. In GSE78220, samples were pair-end sequenced with a read length of 2 × 100 bp (Illumina HiSeq2000) and amplified to the UCSC hg19 reference genome using Tophat2 ^34^ and normalized expression levels of genes were expressed in FPKM values as generated by cuffquant and cuttnorm ^35^. In GSE91061, raw FASTQ files were aligned to the hg19 genome using the STAR aligner (STAR, RRID: SCR_004463), counted, and annotated using Rsamtools v3.2, and the TxDb.Hsapiens.UCSC.hg19.known gene transcript database. Raw reads were normalized using a regularized logarithm transformation function with a robust estimation. The tumor immune syngeneic mouse (TISMO) database was similarly accessed ^36^.

### Statistical Analysis

Descriptive statistics were used to summarize the data. Frequencies and counts were used to describe categorical variables, whereas means, standard deviations, medians, and interquartile ranges were used for numeric data. Chi-square or Fisher’s exact test, as appropriate, was used to compare categorical variables, and the Mann-Whitney test was used for numeric variables. Overall survival was defined as the time from the start of randomization of treatment assignment to the date of sacrifice, as directed by the IACUC guidelines. Hazard ratios and associated 95% confidence intervals were computed using multivariate Cox regression analyses. Proportional hazards were evaluated using the interaction between the log of time and each variable; we found no statistical evidence to indicate proportional hazard violation, given that the P-value was > 0.05. Two-sided p-values less than 0.05 were considered statistically significant. All experimental data was analyzed using Prism 7(GraphPad Prism, RRID: SCR_002798) and SAS version 9.4.

## Results

### High βAlt correlates with ICB response

To test the hypothesis that βAlt would predict response to ICB, we used the IMvigor210 trial dataset in which patients with metastatic bladder cancer were treated with anti-PD-L1 (atezolizumab) and platinum chemotherapy ^10,31^. Patients were classified according to RECIST criteria as responders, which encompassed complete response and partial response, and as non-responders if the disease remained stable or progressed. βAlt was assessed by dimensionally collapsing the correlation of a chronic TGFβ signature consisting of 50 TGFβ target genes and a signature composed of 36 alt-EJ repair genes that TGFβ suppresses ^23, 24^ (*Supplemental Table 3)*. A high βAlt score represents cancers in which TGFβ signaling is low and error-prone DNA repair is high.

As previously reported for TCGA and other datasets, the anti-correlation between the expression of these signatures, indicative of their functional relationship in which TGFβ suppresses alt-EJ gene expression was recapitulated in the unsupervised clustering of RNA-seq data (n=348) from IMvigor210 specimen (**Fig. 1A**). Single-specimen gene set enrichment (ssGSEA) of both signatures were also significantly anti-correlated (Pearson Correlation Coefficient (PCC) r= -0.48, *P<*0.0001; **Fig. 1B**). Next, we analyzed βAlt as a function of ICB response. The mean βAlt was significantly (*P<*0.0001) larger in responders (R, n=68, x̄=17.5) than non-responders (NR, n=230;x̄ = -10.1; **Fig. 1C**). To validate this finding, we analyzed the GSE78220 dataset reported by Hugo et al. from metastatic melanoma patients treated with anti-PD-1 (pembrolizumab), in which 15 of 27 (55%) patients were classified as responders ^32^. As with IMvigor210, the βAlt scores of responders were significantly greater (*P<*0.05, Mann-Whitney test, 95% CI {-0.42, -63.0}) compared with those whose disease progressed (*Supplemental Fig. S1A*).

**Fig. 1.**
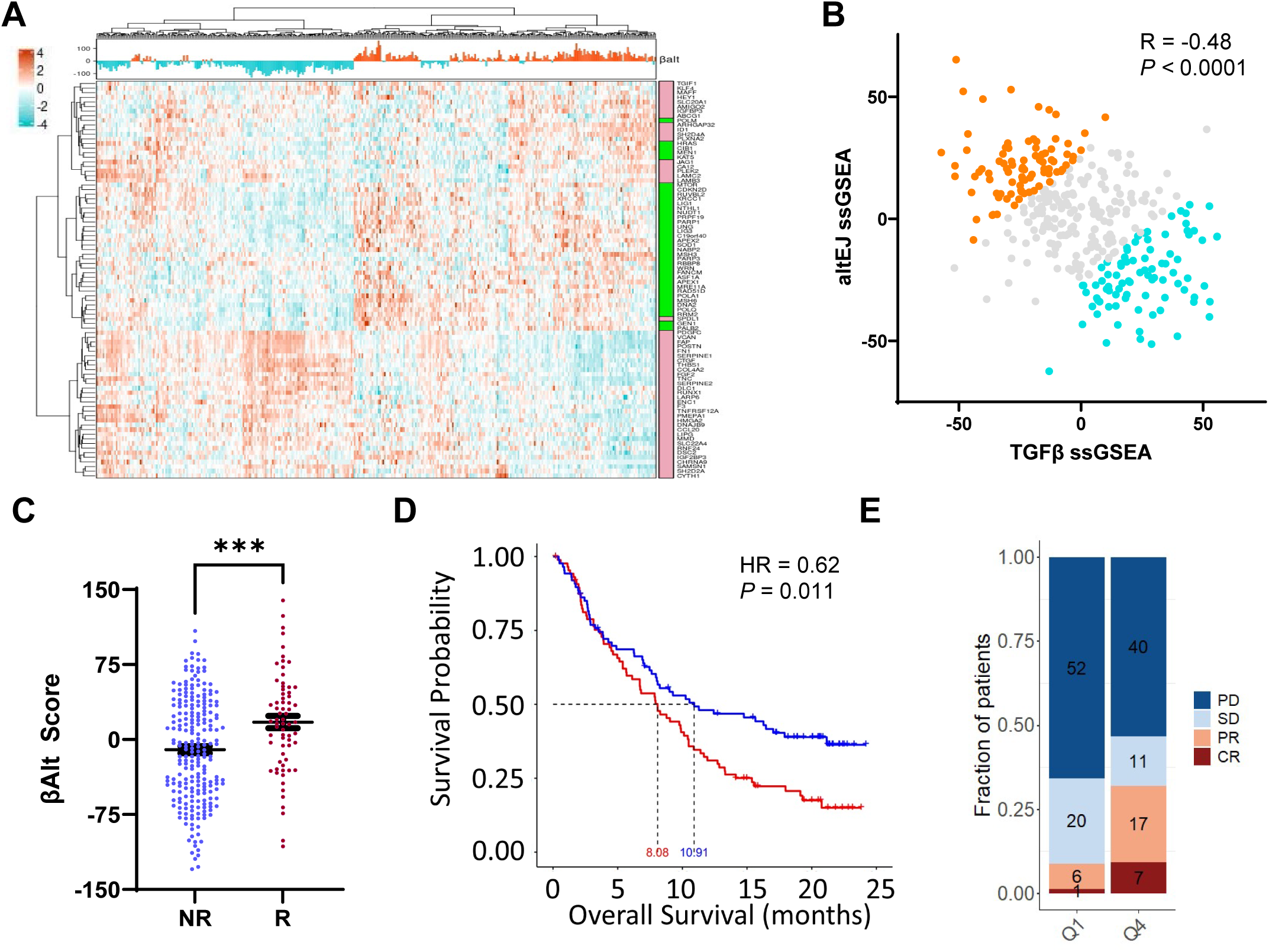
High βAlt is associated with ICB response. Analysis of data from IMvigor210 metastatic bladder patient (n=348). **(A)** Unsupervised hierarchical clustering heatmap depicting TGFβ (pink) and alt-EJ (green) gene signatures of patients (n=348). βAlt score is annotated above (coral=high, teal=low). **(B)** TGFβ and alt-EJ ssGSEA scores are negatively correlated (R=-0.48, P<0.0001). The high quartile (Q4; n=87, blue) and low quartile (Q1, n=87, red) are indicated. **(C)** The IMVigor210 patients defined as responders (R, n=68) have significantly higher mean βAlt score oxf ( = 17.5 ± 6.1 SEM) versus those designated non-responders (NR, n=230; x= -10.1 ± 3.4 SEM), P<0.0001, Wilcoxon test. **(D)** Overall survival of patients in high (Q4, blue) versus low (Q1, red) βAlt quartiles designated in 1B. Kaplan-Meier analysis, HR=0.62, P=0.011 calculated via log-rank test. **(E)** Distribution of response types (complete response, CR; partial response, PR; progressive disease, PD; stable disease, SD) for βAlt Q1 (n=79) and Q4 (n=75). Q4 is significantly enriched for CR and PR. P=0.0031, Chi-square test.

Patients whose tumors fell in the high βAlt quartile (Q4) had a median survival of 10.9 months and experienced better overall survival (hazard ratio 0.62, *P=*0.011) compared with those ranked as low βAlt levels (Q1), whose median survival was 8.1 months (**Fig. 1D**). The fraction of patients who were responders was significantly enriched in Q4 versus Q1 (*P* =0.0031; **Fig. 1E**). Indeed, two-thirds of the IMvigor 210 responders had βAlt scores above 0 (45/68, 66%). IMvigor210 comprised two cohorts of patients: cohort 1 was platinum ineligible and treatment naïve at the time of immunotherapy, cohort 2 had failed prior treatment with platinum chemotherapy ^37, 38^. Notably, the frequency of complete and partial response by RECIST criteria was significantly greater for βAlt quartile 4 compared with quartile 1 in both (Cohort 1 P=0.0045; Cohort 2 P=0.023; *Supplemental Fig. S1B*).

Mariathasan et al. ^10^ reported that patient response was significantly associated with tumor mutation burden. As expected from error-prone DNA repair, the βAlt score is also associated with the fraction of genome altered ^23^. βAlt is significantly correlated with tumor mutation burden, as well as predicted neoantigen frequency, in IMvigor210(*Supplemental Fig. S1C,D*). Mariathasan et al. also reported that a TGFβ expression signature is associated with the immune excluded pattern of lymphocyte distribution as a mechanism of resistance. Clustering of patient data based on Mariathasan et al assignment of immune infiltrate and annotated for prior chemotherapy, response, βAlt score, and TMB revealed that the range of high βAlt score is not exclusive to a particular pattern of lymphocyte distribution (*Supplemental Fig. S1E*).

The significant association of βAlt with response to ICB in two patient populations substantiates the predictive value of high βAlt, which also reports sensitivity to genotoxic treatments due to poor DNA repair ^23, 24^.

### High βAlt correlates with the immune composition in a tissue-agnostic manner

Next, we investigated the specific immune context based on the expectation that high βAlt, like dMMR tumors ^17^, would be infiltrated. The challenges of classifying the frequency and pattern of tumor-infiltrating lymphocytes and immune composition objectively and reproducibly have prompted several approaches ^39^. Here, we used 10 tumor-educated immune signatures (TeIS; *Supplemental Table 4*) tumor defined by immune profiling immune cells isolated from 364 surgical tumor specimens across 12 tissues ^29^. We conducted unsupervised clustering of TeIS expression signatures for IMvigor210 specimens that resulted in an expression gradient from immune-rich to immune-poor (**Fig. 2A**). Annotation of this heatmap for βAlt showed a broad parallel transition from immune-rich, low βAlt to immune-poor, high βAlt. Separating the cluster with the highest and the lowest mean TeIS expression confirmed that immune-poor tumors had significantly higher βAlt scores when compared with immune-rich tumors (*P<*0.001, **Fig. 2B**). Of the IMvigor 210 responders with the highest βAlt scores (31/68), 74% were classified as immune-poor (23/31). Consistent with an immune cold phenotype, the immune-poor group had significantly lower expression of a type I IFN gene signature ^40^ (**Fig. 2C**, *Supplemental Fig. S2A*). Assessing TeIS as a function of lymphocytic distribution assigned in Mariathasan et al., showed that desert tumors were enriched for immune-poor TeIS signatures, inflamed tumors were predominantly immune-rich, and excluded tumors exhibit a range of TeIS scores (*Supplemental Fig. S2B*).

**Fig. 2.**
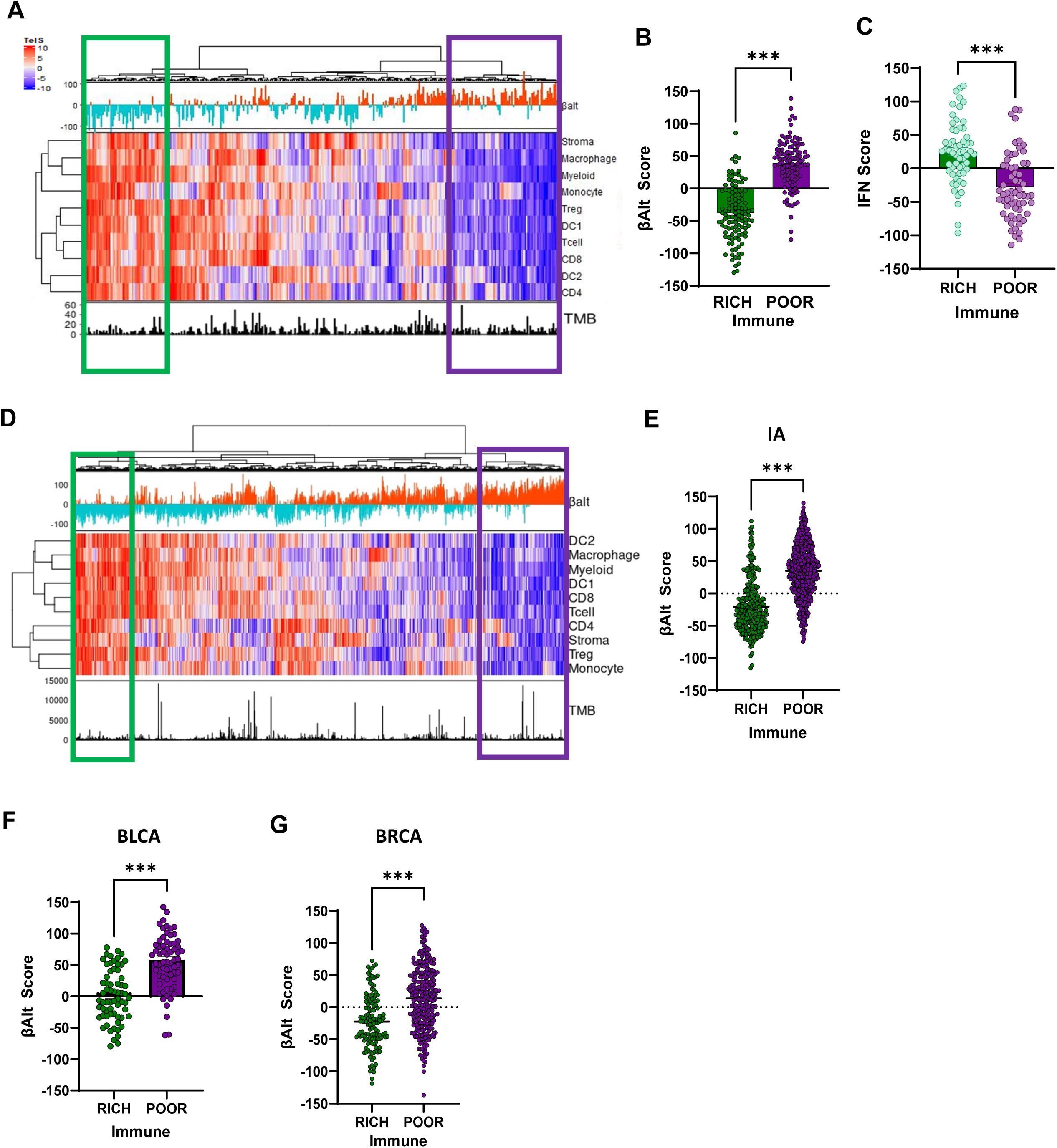
High βAlt correlates with the immune composition in a tissue-agnostic manner. **(A)** Unsupervised clustering of TeIS heatmap IMvigor210 metastatic bladder patient (n=348) annotated for βAlt score above and tumor mutation burden (TMB) below. The green box designates an immune-rich cluster (n=116); the purple box designates an immune-poor cluster (n=116). **(B)** βAlt scores of IMvigor210 tumors classified as immune-rich are significantly lower than those designated as immune-poor (*P<*0.001; Wilcoxon test). **(C)** IFN signature scores of ImVigor 210 immune-poor tumors are significantly lower than immune-rich tumors. (*P<*0.001; Wilcoxon test). **(D)** Unsupervised hierarchical clustering of TeIS signatures for immune archetype TCGA dataset (n=4341) from Combes et al. 2021 annotated with βAlt scores (red=positive, teal=negative) above and tumor mutation burden (TMB) below. The immune-rich cluster is designated by green box, the immune-poor cluster is designated by a purple box. **(E-F)** βAlt score distribution of immune-rich (green) quartile versus immune-poor (purple) quartile for: **(E)** TCGA IA data set from Combes et al; immune-rich (n=297) and immune-poor (n=523). Wilcoxon test. **(F)** TCGA bladder cancer (BLCA; immune-rich, n=65; immune-poor, n=67). Unpaired T test. **(G)** TCGA breast cancer (BRCA; immune-rich, n=135 and immune-poor, n=267). Unpaired T test. Statistical significance denoted as \*\**P<*0.01, *\*\*\* P<*0.001.

The unexpected characterization of high βAlt as immune-poor tumors led us to test the association of βAlt across a subset of TCGA solid tumors (n=4341) that had been assigned immune archetypes ^29^. Unsupervised clustering of TeIS replicated the parallel transition of low to high βAlt with immune-rich to immune-poor (**Fig. 2D**) observed in IMvigor210 bladder cancer. Likewise, immune-poor cancers had the highest βAlt scores (**Fig. 2E**). We then examined the entire TCGA bladder cancer cohort, which also showed a similar parallel transition displayed as a heatmap (*Supplemental Fig.S2C);* immune-poor tumors had significantly higher βAlt scores (**Fig. 2F**). To ascertain whether this relationship was generalizable, we analyzed breast cancer that were not included in the immune archetype. The TCGA breast cancer specimens showed a similar pattern of immune-rich to poor TeIS, paralleled by low to high βAlt (*Supplemental Fig. S2D*); immune-poor tumors had significantly higher βAlt scores (**Fig. 2G**). Consistent with the correlation of high βAlt with immune-poor tumors, the expression of markers of cytotoxic T cells was uniformly negatively correlated with βAlt across cancer types (*Supplemental Fig. S2E*). Given the compelling evidence that dMMR, which has high predictive value in immunotherapy, gives rise to inflamed tumors ^41^, it was particularly surprising that genomically unstable ^23^ and radiosensitive ^24^ high βAlt cancers are immune-poor and IFN low.

### High βAlt, immune-poor tumors can convert to immune-rich with treatment

It is well-established that loss of cell-intrinsic TGFβ signaling in cancer cells can lead to increased TGFβ activity, which was elegantly demonstrated in the KRAS lung model using tumor-specific CRISPR knockdown of *TGBR2* that resulted in lung tumors devoid of lymphocytes ^42^. Hence, loss of TGFβ signaling could establish a highly immunosuppressive tumor microenvironment (TME) but how such tumors respond to ICB is unknown. To investigate this experimentally, we determined whether the association of βAlt with the immune context was evident in mouse tumors.

The tumor immune syngeneic mouse (TISMO) database consists of 1500 mouse tumors covering 68 cell lines and 19 cancer types ^36^. Unsupervised hierarchical clustering of untreated specimens using the TeIS signatures also resulted in a gradient from immune-rich to immune-poor accompanied by increasing βAlt scores (*Supplemental Fig. 3A,B*). Twelve independently derived mammary tumor-derived transplants (mTDT) also exhibited this relationship (*Supplemental Fig. 3C,D*). Nearly half of the high βAlt mouse tumors were immune-poor (47%, 11/23). Family F mTDT clustered in the far right arm of the dendrogram with the higest βAlt score and poorest TeIS expression. The recapitulation in mouse models of the relationship between βAlt and immune context underscores its high conservation and provided the means to investigate how to therapeutically exploit it.

To this end, we used a high βAlt, low TeIS mTDT (family F) to investigate the response to TGFβ inhibitors with an anti-PD-L1 ^27^. Even though high βAlt cancer cells are unresponsive to TGFβ, we hypothesized that TGFβ inhibition (TGFβi) would be necessary to release immunosuppression. To reveal the immunogenic potential of alt-EJ misrepair, we incorporated radiation into our treatment strategy considering that high βAlt tumors are sensitive to radiation, which could provoke antigen release ^43^. Mice bearing mTDT with an average volume of approximately 75 mm^3^ were randomized into treatment arms (**Fig. 3A**), consisting of 10 Gy single dose radiation (RT), anti-PD-L1 or TGFβi, dual combinations, or triple treatment. Monotherapy with anti-PD-L1 or TGFβi, as well as their combination, had little effect on survival. In contrast, triple treatment that included RT led to a significant increase in overall survival (**Fig. 3B**). The median survival time for monotherapy consisting of anti-PD-L1 or TGFβi and their dual combination was 8 days, which increased to 11 days with RT only. Dual therapy with RT plus anti-PD-L1 or TGFβi led to a median survival of 13 days, whereas triple treatment (combination plus RT) increased the median survival to 22 days.

**Fig. 3.**
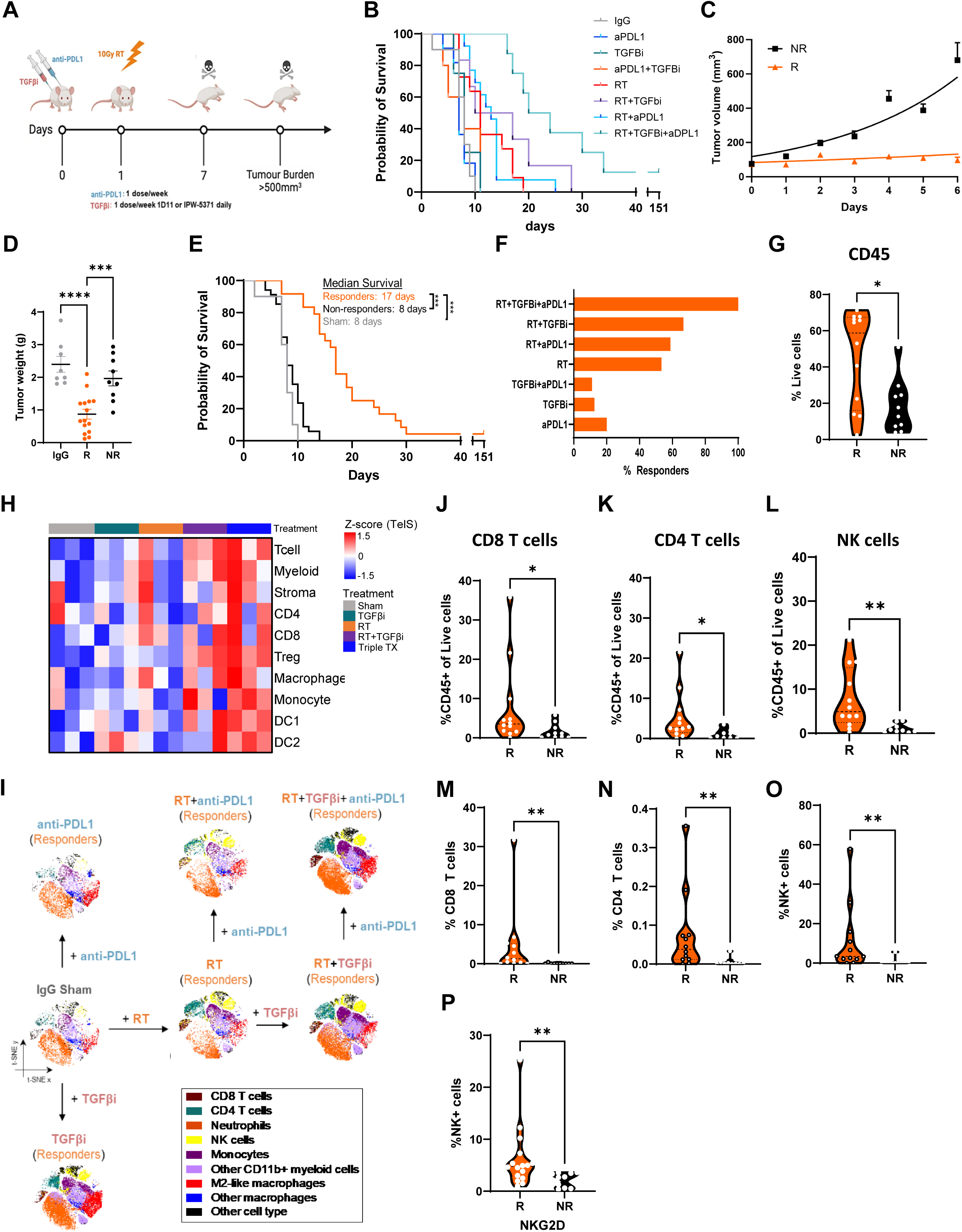
High βAlt, immune-poor tumors respond to ICB by converting to immune-rich. **(A)** Experimental schematic of treatment schedule, primary endpoints and tissue collection. **(B)** Kaplan-Meier overall survival curves of mTDT treated with IgG isotype (grey, n=8), anti-PD-L1 (blue, n=7), TGFβi (green, n=8), anti-PDL1+TGFβi (orange, n=8), 10 Gy RT (red, n=11), RT+anti-PDL1 (light blue, n=13), RT+TGFβi (purple, n=6) and RT+TGFβi+anti-PDL1 (teal, n=7). *P<*0.0001 via Log rank test. **(C)** Mice classified as responders (R, orange, n=34) at 7 days post-treatment exhibit significantly decreased (*P=*0.0023; two-way ANOVA) growth trajectories (mean ± SEM) compared with non-responders (NR, black, n=24). **(D)** Tumor weights (mean ± SEM) of mice designated as R (n=15) were significantly less (*P<*0.001, one-way ANOVA) than those from mice treated with IgG isotype (n=8) or those designated as NR (n=9). **(E)** Kaplan-Meier overall survival curves of R (n=24) were significantly longer (*P<*0.001, Log-rank test) than mice designated as NR (n=34) or sham treated (n=10). **(F)** Percentage of responders among treatment groups. **(G)** Percentage of live intra-tumoral CD45+ cells was significantly increased (P<0.05) in tumors designated as R (n=12) versus NR (n=10). **(H)** Heatmap showing TeIS-RNA deconvoluted immune cell populations of mTDT treated with IgG isotype, 10 Gy RT, RT+TGFβi and RT+TGFβi+anti-PDL1 (n=3 per group) 7 days post-treatment. **(I)** t-SNEs of tumor immune cell composition between R treated with same treatments in 3B. **(J-L)** CD45-normalized percentages of live intra-tumoral **(J)** CD8 T cells, **(K)** CD4 T cells and **(L)** NK cells, Proliferation, indicative of activation, marked by Ki67 in **(M)** Ki67+ CD8 T cells, **(N)** Ki67+ CD4 T cells, and **(O)** Ki67+ NK cells was significantly increased in R. **(P)** Percentage of NK cell activation marker NKG2D positive cells was also significantly increased in R. Statistical significance is denoted as \**P<*0.05, ** *P<*0.01, *\*\*\*P<*0.001.

Individual mice manifested two distinct patterns of early responses to treatment: tumors either grew exponentially after treatment or their growth was controlled during the first 7 days post-treatment (*Supplemental Fig. 3E*). This difference, early control versus growth, led us to classify mice across treatment groups as responders and non-responders 7 days post-treatment. A similar classification was recently used to characterize the mouse tumor response to a bifunctional TGFβ trap and anti-PD-1 ^44^. In this fashion, the tumor growth rates showed a highly significant difference (**Fig. 3C**). The doubling time of tumors classified as responders (n=34) was 9 days, and that of non-responders (n=24) was 2.6 days (95% CI {2.2-3.1}). Tumor weight was also significantly lower in responders than in non-responders (Mann-Whitney test, *P=*0.007, 95% CI {-0.42, -1.76}, **Fig. 3D**). Importantly, the overall survival of mice classified as responders was significantly better than that of non-responders (**Fig. 3E**). The median survival was 8 days for untreated mice or non-responders. In contrast, it was more than doubled to 17 days in those classified as responders (log-rank, *P<*0.0001).

The proportion of responders paralleled the survival outcomes with regard to therapy modality (*P=*0.0023, two-way ANOVA, **Fig. 3F**). Approximately 15-20% of the mice treated with either anti-PD-L1 or TGFβi as a monotherapy or dual combination were responders. The proportion of responders to RT tripled to 53%. In combination with RT, anti-PD-L1 therapy had little effect (58%), whereas RT and TGFβi increased the proportion of responders to 67%. All mice (9/9) were classified as responders following triple treatment with RT, anti-PD-L1, and TGFβi.

Uncoupling population responses from individual responses allowed the investigation of biological correlates of response to pinpoint early critical differences between responders and non-responders (*Supplemental Fig. 3F*). The frequency of CD45+ leukocytes was significantly higher in tumors classified as responders than in those compared with non-responders (Mann-Whitney test, *P=*0.02, **Fig. 3G**). Consistent with this, analysis of bulk RNA-sequencing data from treated tumors at 7 days using TeIS showed increased intra-tumoral immune cell infiltration only when radiation was combined with TGFβi, which was amplified with the addition of anti-PD-L1, essentially shifting the immune cold tumor to hot **(Fig. 3H)**. This conclusion led to further functional investigation via multispectral flow cytometry. To identify and visualize immune cell populations, unsupervised analysis was performed using the dimensionality reduction algorithm t-distributed stochastic neighbor embedding (t-SNE) of responders across treatment groups (**Fig. 3I**). Plots from individual monotherapy treatments implicate RT and TGFβi as drivers of compositional changes in responders, whereas anti-PD-L1 had little effect. The frequency of tumor-infiltrating CD8+ and CD4+ T cells increased in responders (**Fig. 3J-K**). The most prominent cell type increased in responders was natural killer (NK) cells (Mann-Whitney test, *P=*0.003, 95% CI {-1.22, -13.3}, **Fig. 3L**). Consistent with activation and clonal expansion, the proliferation of CD4+ and CD8+ T cells (Mann-Whitney test, *P=*0.003 **Fig. 3M-N**) and NK cells (Mann-Whitney test, *P=*0.002; **Fig. 3O**) was significantly increased in responders. In contrast, neither a number nor proliferation increase was evident in non-responders, regardless of treatment (*Supplemental Fig. 3G, H*). Moreover, increased immune reactivity within the tumor was paralleled by increased CD8+, CD4+ T cells, and NK cells in the blood of responders compared with non-responders (*Supplemental Fig. 3I-K*) whereas other blood immune cells did not change. The percentage of NK cells specifically increased in responders (**Fig. 3L)**, which also exhibited, NKG2D, a marker of NK cell activation (Mann-Whitney test, *P<*0.001; **Fig. 3P**). Hence, triple treatment converted a high βAlt, immune-poor tumor to immune-rich accompanied by significant activation of immune cells.

### NK cells mediate the response of βAlt high, immune-poor tumors to ICB therapy

NK cells were significantly increased in responders compared with non-responders or sham (ANOVA, *P<*0.008; *Supplemental Fig. 4A*). The proportion of NK cells among responders (n=13) nearly tripled from 2.8% in non-responders (n=10) to 8% in responders. Moreover, activation, as indicated by Ki67 proliferation, markedly increased (*Supplemental Fig. 4B*). Indeed, treatment with TGFβi in combination with RT resulted in the most significant expansion and activation of NK cells compared with RT alone or RT combined with anti-PD-L1 (**Fig. 4A-C**). These findings were exemplified by the significant enrichment of NK cell activation signatures present in the transcriptomics from tumors treated with RT and TGFβi compared with controls **(Fig. 4D).**

**Fig. 4.**
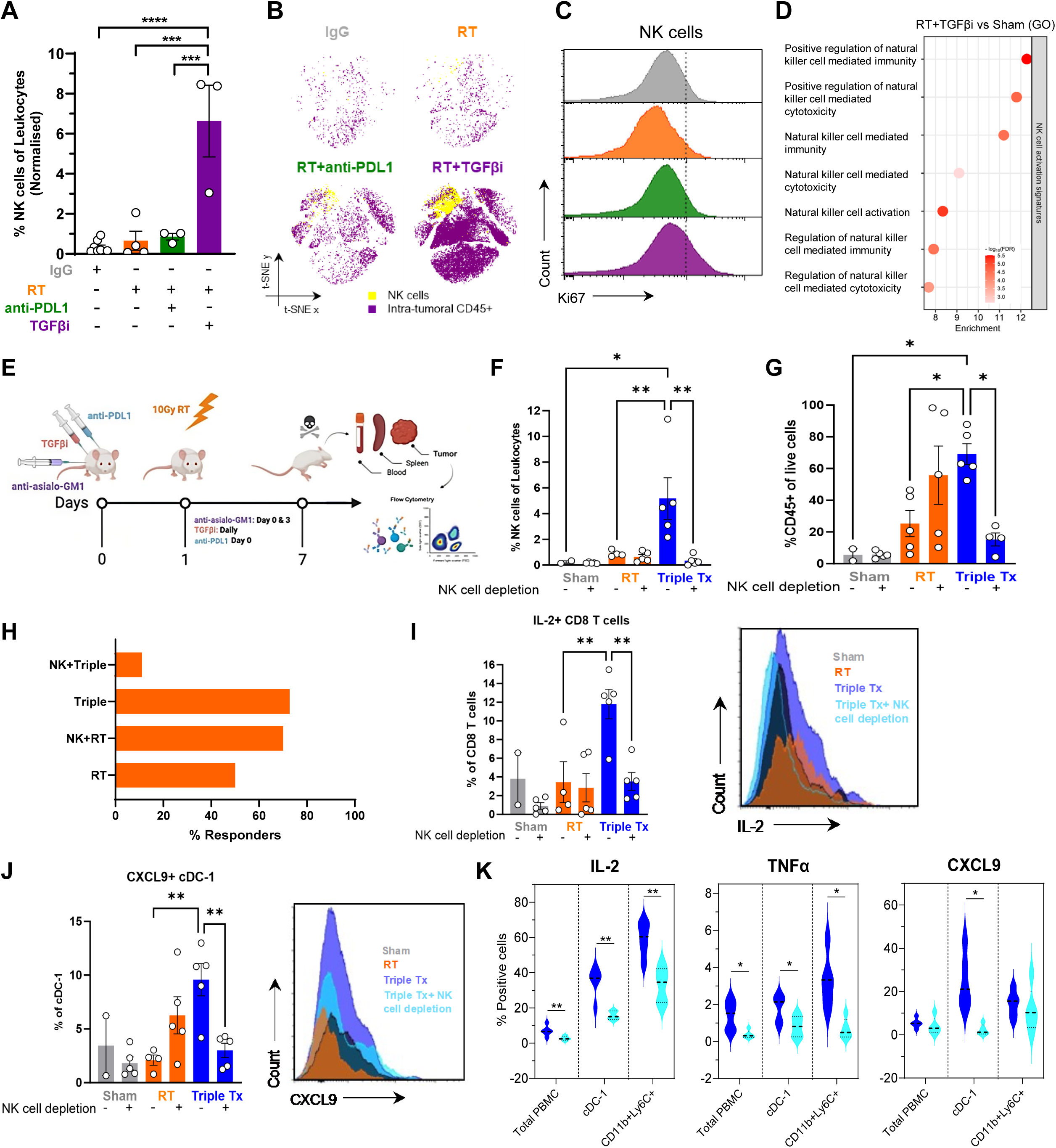
NK cells mediate the response of βAlt high, immune-poor tumors to ICB therapy. **(A)** CD45%-normalized percentage of NK cells in mTDT treated with RT + TGFβi (purple, n=3) were significantly increased compared with IgG (grey, n=8), RT (orange, n=4), RT + anti-PD-L1 (green, n=3). One-way ANOVA. **(B)** t-SNEs depicting tumor NK cells (yellow) and CD45+ (purple) cells in mTDT treated as above. Depicted cell numbers across treatments have been normalized for CD45+ infiltration. **(C)** Representative fluorescence histograms of Ki67 in tumor-infiltrating NK cells treated as in A. **(D)** Gene ontology enrichment for NK cell activation signatures in mice treated with RT + TGFβi compared with sham-treated 7 days post-treatment. **(E)** NK cell depletion experimental schematic of treatment schedule, primary endpoint, and tissue collection. Percentages of live intra-tumoral **(F)** CD45+ cells and (G) %CD45-normalized NK cells. **(H)** Percentage of responders between treatment groups. Percentages of live intra-tumoral **(I)** %CD8 T cells IL-2+ and **(J)** %cDC-1 C XCL9+ cells between F mTDT treated with IgG isotype without (n=2) or with NK cell depletion (n=5), RT without (n=5) or with (n=5) NK cell depletion and RT+TGFβi+anti-PDL1 without (n=5) or with (n=5) NK cell depletion, representative fluorescence histograms (right). **(K)** Percentage of circulating IL-2+, TNFα, and CXCL9 in total PBMC (left), cDC-1 (middle) and CD11b+Ly6C+ cells (right) increased in mTDT following triple TX (blue) versus triple TX + NK depletion (light blue). Unpaired t-tests. Data are depicted as mean ± SEM; each point represents a mouse, and one-way ANOVA P-values are represented as follows: * *P<*0.05, ** *P<*0.01, *** *P<*0.001.

We postulated that the increase in NK cells was the key to the recruitment of CD4 and CD8 T cells and the response to anti-PD-L1. High βAlt tumors are intrinsically deficient in DNA repair. Recent studies have shown that blocking DNA repair in irradiated tumors can boost the response to ICB therapy by activating NK cells ^45^. Moreover, TGFβ broadly downregulates chemokine receptor expression necessary to recruit NK cells to the tumor and impairs NK cell function ^46^.

To test whether NK cells were critical, a NK depleting antibody was administered before RT alone or triple treatment with TGFβi and anti-PD-L1 (**Fig. 4E**). The efficacy of depletion was confirmed by the analysis of PBMC, in which both NK cells and proliferating NK cells were significantly decreased (*Supplemental Fig. 4C, D*). As in the prior study, infiltrating leukocytes increased significantly in tumors treated with the triple combination over RT alone, which was reduced to that of untreated tumors by NK depletion (**Fig. 4F**). Intratumoral NK cells, which were negative for a marker of innate lymphoid cell 1 (*Supplemental Fig. 4E*), were depleted in the triple treatment arm to negligible levels of the untreated tumors (**Fig. 4G**). NK depletion before triple treatment (n=12) completely abrogated the 75% response rate of this combination (Chi-square, *P<*0.0001) (**Fig. 4H**). The response to RT significantly increased (Chi-square, *P=*0.0004) from 50% (n=11) to 70% (n=11).

NK cells are controlled through specific cell-surface receptors that can either stimulate or inhibit the activity of NK cells, and their combinatorial signaling dictates NK cell response and function ^47^. The balance of activating to inhibitory signaling conditions in the NK cell-target synapse can be shifted by cytokines, such as TGFβ. We examined intracellular cytokine levels to understand better the mechanisms mediating the NK cell axis in high βAlt immune-poor tumors. Although CD8 T cells were not increased across treatments, IL-2+ CD8+ T cells were significantly increased in triple-treated tumors compared with untreated or irradiated tumors, which was abrogated by NK cell depletion (**Fig. 4I**). Similarly, classical dendritic cell (cDC) numbers were similar across treatments, but CXCL9+ cDCs were significantly increased only in tumors following triple combination treatment (**Fig. 4J**). We hypothesize that the crosstalk of activated NK cells and cDC maintains a positive feedback loop for chemotactic recruitment of immune cells through CXCL9 release. Levels of the cytokines IL-2, TNFα, and CXCL9 in circulating immune cells, which were increased by triple treatment, were also NK cell-dependent (**Fig. 4K)**. Thus, the unique TME of high βAlt tumors can be primed for checkpoint blockade by promoting an NK cell-dependent response, detectable both in the tumor and circulation.

### High βAlt, immune-poor tumors convert to immune-rich that respond to ICB

Restoration of NK cell function in cancer has been hypothesized to activate tumor immunity ^47^. To determine the generalizability of the mechanism, we used the TISMO database, which comprises data from syngeneic mouse tumors treated with ICB. We identified two widely used tumor models, melanoma B16 and colorectal CT26, which are high βAlt and immune-poor, as measured by TeIS at baseline (**Fig. 5A, C**). Notably, treatment with ICB shifted most of the tumors to an immune-rich phenotype (*P<*0.0001, chi-square test). Moreover, the gene ontology signatures of NK activation were significantly enriched in responders (**Fig. 5B, D**).

**Fig. 5.**
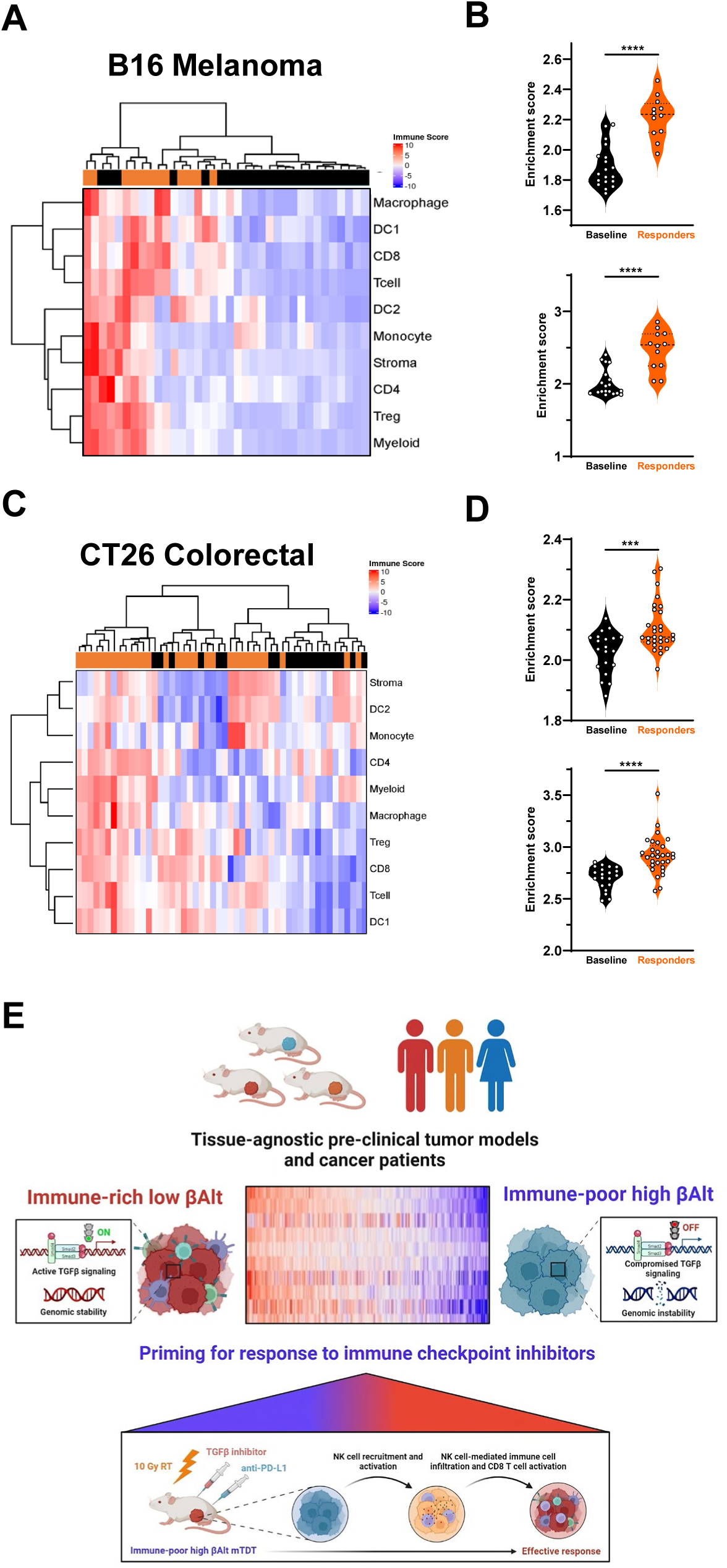
High βAlt, immune-poor tumors convert to immune-rich upon response to ICB. **(A)** Unsupervised hierarchical TeIS clustering of B16 melanoma model at baseline (black, n=19) versus responsive to ICB treatment (orange, n=12). **(B)** Gene ontology signatures of NK activation (top) and NK mediated immunity (bottom) of B16 baseline versus response to ICB treatment. Unpaired T test. **(C)** Unsupervised hierarchical TeIS clustering of CT26 melanoma model at baseline (black, n=20) versus responsive to ICB treatment (orange, n=30). **(D)** Gene ontology signatures of NK activation (top) and NK mediated immunity (bottom) of CT26 baseline versus response to dual ICB treatment. Unpaired T test. **(E)** Graphical summary depicting low βAlt tumors, which have active TGFβ signaling and genomic stability, are immune-rich, while high βAlt tumors that have low TGFβ signaling and genomic instability are immune-poor. Both human data and experimental studies indicate that NK cell-mediated lymphocyte recruitment facilitates ICB response in high βAlt, immune-poor tumors.

To ascertain whether this mechanism is evident in human cancers, we analyzed the GSE91061 dataset in which paired biopsies from advanced melanoma patients (n=43) were obtained before and after treatment with anti-PD-1 (nivolumab) ^33^. Nine patients were classified as responders to anti-PD-1 therapy and 34 as non-responders. As with IMvigor210, TGFβ and alt-EJ signatures were anti-correlated, although there was no significant difference in the mean βAlt of complete and partial responders compared with stable or progressive disease pre-treatment in this small group. Based on TeIS unsupervised clustering, there was however a marked association between high βAlt levels and immune-poor TME (*Supplemental Fig. 4F*). We compared the pretreatment and on-treatment TeIS signatures of responders for whom transcriptomics was available (n=9). Two responders, classified as high βAlt and immune-poor pre-treatment, moved to immune-rich in the on-treatment biopsy (*Supplemental Fig. 4G*). None of the patients classified as immune-rich moved on treatment, regardless of the βAlt score. Genes associated with NK activation also increased in high βAlt paired biopsy melanoma patients in response to anti-PD-1 therapy (*Supplemental Fig. 4H*). Although the sample size is small, these human data support the experimental studies whereby NK cell-mediated lymphocyte recruitment facilitates ICB response in high βAlt, immune-poor tumors. We suggest that βAlt and TeIS provide the means to prospectively identify a previously unrecognized subset of immune-poor patients in whom TGFβ inhibition reverses resistance to ICB (**Fig. 5E**).

## DISCUSSION

Predicting patient response to immunotherapy is complicated by an incomplete understanding of the immune context ^48^, the diversity of the systemic immune environment ^49^, and the complexity of the drivers of cancer immunity ^50^. Here we show that patients with high βAlt tumors, which have compromised TGFβ signaling and DNA repair, are immune-poor yet responsive to ICB treatment. We hypothesized that loss of feedback resulted in compensatory TGFβ generation that was masking the immunogenic potential of alt-EJ misrepair. Preclinical modeling identified that a therapeutic combination of TGFβ inhibition and radiation was able to instigate conversion from immune-poor to immune-rich, evidenced by lymphocytic infiltration. Analysis of high βAlt, immune cold mouse tumors and clinical trial data showed that NK cell activation was instrumental in converting immune-poor tumors to immune-rich. These data identifying ICB-responsive cold cancers in which intrinsic loss of TGFβ signaling generates TGFβ rich, immunosuppressive TME suggest that the use of βAlt scoring as a biomarker to selectively deploy TGFβ inhibitors in immunotherapy.

The precise cellular and molecular pathways that lead some patients to respond, while others do not, remain largely undefined because the complex biology of the immune microenvironment is incompletely understood ^50^. Although lymphocytic infiltration is generally associated with response, there needs to be a sufficient and accurate predictor, as some patients with poorly infiltrated tumors still benefit from ICB. In contrast to the inflamed phenotype of dMMR ^13, 51^, the genomic and phenotypic consequences of compromised TGFβ signaling and error-prone alt-EJ repair result in an immune-poor tumor lacking lymphocytes and interferon signaling. The paradoxical consequence of high βAlt is an immunosuppressive TME, underscoring the tightly coupled intrinsic and extrinsic actions of TGFβ. Whereas the DNA damage from dMMR elicits type-I IFN that recruits lymphocytes, the high levels of damage from alt-EJ are overridden by an established TGFβ-rich TME. We speculate that cancer cells that lose TGFβ signaling competency early in carcinogenesis increase TGFβ activity, which establishes an immunosuppressive tumor microenvironment. It may be that loss of TGFβ also compromises innate DNA sensing mechanisms, thereby indirectly suppressing IFN, further endorsing the immune cold TME.

The clinical success of cancer immunotherapy is predicated on disrupting the equilibrium established between the patient’s cancer and their immune system ^52^. A deeper understanding of the diversity of the systemic immune environment across human malignancies is critical for improving immunotherapy treatment strategies. Our analysis was aided by the immune archetype paradigm based on the immune profiling initiative, which conducts a holistic survey across cancers ^29^. These analyses and preclinical data show that high βAlt, immune-poor mouse, and human tumors represent a category of malignancies that can be shifted from immune-poor to immune-rich by different treatments and thus have the potential to benefit from ICB. We found that radiation and TGFβ inhibition were able to activate T cells via NK cells in the high βAlt, immune-poor mTDT. The widely used high βAlt and immune-poor B16 melanoma and CT26 colorectal models also shift to immune-rich in response to ICB. Although there are limited data from humans, a small, paired biopsy study reported by Riaz et al. ^33^ shows that high βAlt, immune-poor melanoma cancers can shift to immune-rich in response to ICB therapy.

We found that TGFβ inhibition leads to activation of NK cells, which are well known to be suppressed by TGFβ ^53–55^, and provide an actionable route by which immune-poor, high βAlt tumors can respond to ICB. In contrast to the prominantj role of CD8 T cells in the ICB response of immune-infiltrated tumors, we found that NK depletion completely abrogated both lymphocytic infiltration and the ICB response in high βAlt and immune-poor mTDT. Moreover, NK cell activation signatures increased upon treatment response in high βAlt, immune-poor F mTDT, B16, and CT26 mouse tumors, and melanoma patients. NK cells correlate with better overall survival in various solid cancers, including gastric, breast, and renal cell carcinoma, and increased response to checkpoint blockade immunotherapy in melanoma ^56, 57^. Immunosuppressive TGFβ broadly downregulates chemokine receptor expression necessary to recruit NK cells to the TME and impairs NK cell function by limiting NK cell antibody-dependent cellular cytotoxicity ^46^. Although the mechanisms mediating the suppression of the NK cell axis are not yet fully understood, the data reported here highlight the necessity of NK cell-mediated immune cell recruitment to convert high βAlt, immune-poor to tumors responsive to ICB.

Because the immune system is a distributed organ that has a regulated repertoire of mechanisms to suppress cancer, research on cancer immunity is moving away from tissue-of-origin specificity toward tissue-agnostic ecosystems and processes ^58, 59^. The more nuanced classification of cancer by immune archetypes is a significant advancement that will enhance patient outcomes by enabling the selection of optimal therapeutic strategies. Although how cancer cells determine the granular immune TME is still unclear ^60^, a high βAlt signature describes a TGFβ incompetent tumor cells embedded in a TME in which TGFβ is the major determinant of immunosuppression.

The main limitation of our study stems from the retrospective analysis of clinical trial data, whereas the prospective application of βAlt to stratify patients for optimal treatment would provide the strongest evidence of clinical utility. We assessed immune context by one classification scheme, immune archetypes, but there are many immune signatures that may provide additional information. Also, there is a growing appreciation of spatial patterning as an important determinant of response to therapy that could add to our assessment of the high βAlt tumor phenotype. In experimental modeling, the high βAlt tumor was derived from breast, whereas the ICB trial data were bladder cancer or melanoma; there may be tissue-specific features that modify ICB response in a manner not captured by our analysis. Indeed breast cancer is one of the least ICB responsive solid cancers ^61^. Our preclinical modeling combined radiotherapy and ICB, yet most trials of such combinations have not shown significant benefit at the population level ^62^. We postulate that radiation-induced cell death reveals the immunogenic potential of DNA misrepaired by alt-EJ, but whether this will be borne out in humans and its potential context-specificity remains to be seen.

To date, no TGFβ inhibitors have been approved in cancer therapy, despite strong preclinical evidence that TGFβ activity drives malignancy and immunosuppression. We suggest that our studies provide the means to prospectively identify a previously unrecognized subset of immune-poor patients in whom TGFβ inhibition reverses resistance to ICB. Further validation of the βAlt signature to predict patient outcomes requires analysis of additional data sets, ideally ones in which paired biopsies and transcriptomics are available pre- and post or during ICB. Substantial opportunity exists to use βAlt and TeIS scores to inform TGFβ inhibitor trials that are underway in combinations with ICB for multiple indications, including therapeutic avenues to activate NK cells and NK cell therapies that might be particularly effective in high βAlt tumors.

## Declarations

## Acknowledgments

The authors would like to thank Drs. David Nyguen and Hang Chang for data analysis, Trevor Jones and Colin Foster, and Hui Zhang in the Barcellos-Hoff lab, and Bushra Samad of the UCSF Data Science Co-Lab for their support of this research. The results presented here are partly based on the analysis of data generated by the TCGA Research Network (https://www.cancer.gov/tcga), and we would like to express our gratitude to the TCGA consortia and their coordinators for the data provision and clinical information used in this study.

## FUNDING

Genentech imCORE network project UCS-3 (MHBH)

Genentech imCORE network project UCS-7 (MHS)

Genentech imCORE network project UCS-12 (MHBH)

National Institutes of Health grant R01CA190980 (MHBH)

National Institutes of Health grant R01CA239235 (MHBH)

National Institutes of Health grant R01CA190980S1 (JM)

National Institutes of Health grant T32CA108462 (JM)

The CyTOF instrument was procured with the support of the National Institutes of Health (grant S10 OD018040).

Tissue processing and immunostaining were performed using the Helen Diller Family Comprehensive Cancer Center Pathology Shared Resource, supported by the National Cancer Institute of the National Institutes of Health under Award Number P30CA082103.

IPW was provided under a material transfer agreement with Innovations Pathway, Inc.

## AUTHOR CONTRIBUTIONS

Guarantor: MHBH

Conceptualization: MHBH, JM

Methodology: MHBH, JM

Statistician: AL

Investigation: JM, JG, IG, KY, AJC,

Visualization: MHBH, JM, JG

Funding acquisition: MHBH, MHS

Project administration: MHBH

Supervision: MHBH

Writing – original draft: MHBH, JM

Writing – review & editing: All

## DATA AND MATERIALS AVAILABILITY

### Code Availability

βAltW code: https://github.com/pujana-lab/Under-review-article

### Data Availability

The public availability of the transcriptomic data generated in this study is pending.

## Abbreviations

ICB: immune checkpoint blockade;
TGFβ: transforming growth factor b;
PD-1/PD-L1: programmed cell death protein one or programmed death-ligand 1;
dMMR: defective mismatch repair;
alt-EJ: alternative end-joining;
IFN: interferon;
TeIS: tumor educated immune signature;
NK cells: natural killer cells;
TME: tumor microenvironment;
TSNE: t-distributed stochastic neighbor embedding;
ssGSEA: single specimen gene set enrichment;
PCC: Pearson’s correlation coefficient;
TGFβi: TGFβ inhibitor;
RT: radiation treatment;
cDC: classical dendritic cell;
mTDT: mammary tumor derived transplants
BRCA: breast cancer;
BLCA: bladder cancer;
IA: immune archetypes

