## Supplemental data for "A conserved subset of cold tumors responsive to immune checkpoint blockade"

***Supplemental Fig. 1.*** **High βAlt is associated with ICB response. (A)** βAlt scores of metastatic melanoma patients (n=27) treated with anti-PD-1 as defined as R (n=15) and NR (n=13) (*P*<0.05, Mann-Whitney). **(B-D)** Analysis of data from IMvigor210 metastatic bladder patient (n=348). **(B)** Distribution of response of IMvigor210 patients in cohort 1 (n=32) and cohort 2 ( n=115) (PD=progressive disease, SD=stable disease, PR=partial response, CR= complete response) as a function of βAlt Q1 and Q4 quartiles. Significant correlation of βAlt with **(C)** TMB (r=0.22, P=0.00026) and **(D)** neoantigens (r=0.22, P=0.0004) . Pearson correlation coefficient, R, t-tailed. **(E)** Clustering of patient data based on Mariathasan et al. assignment of immune infiltrate and annotated for prior chemotherapy, response, βAlt score, and TMB. Note that the βAlt score range is not specific to a particular pattern of lymphocyte distribution.

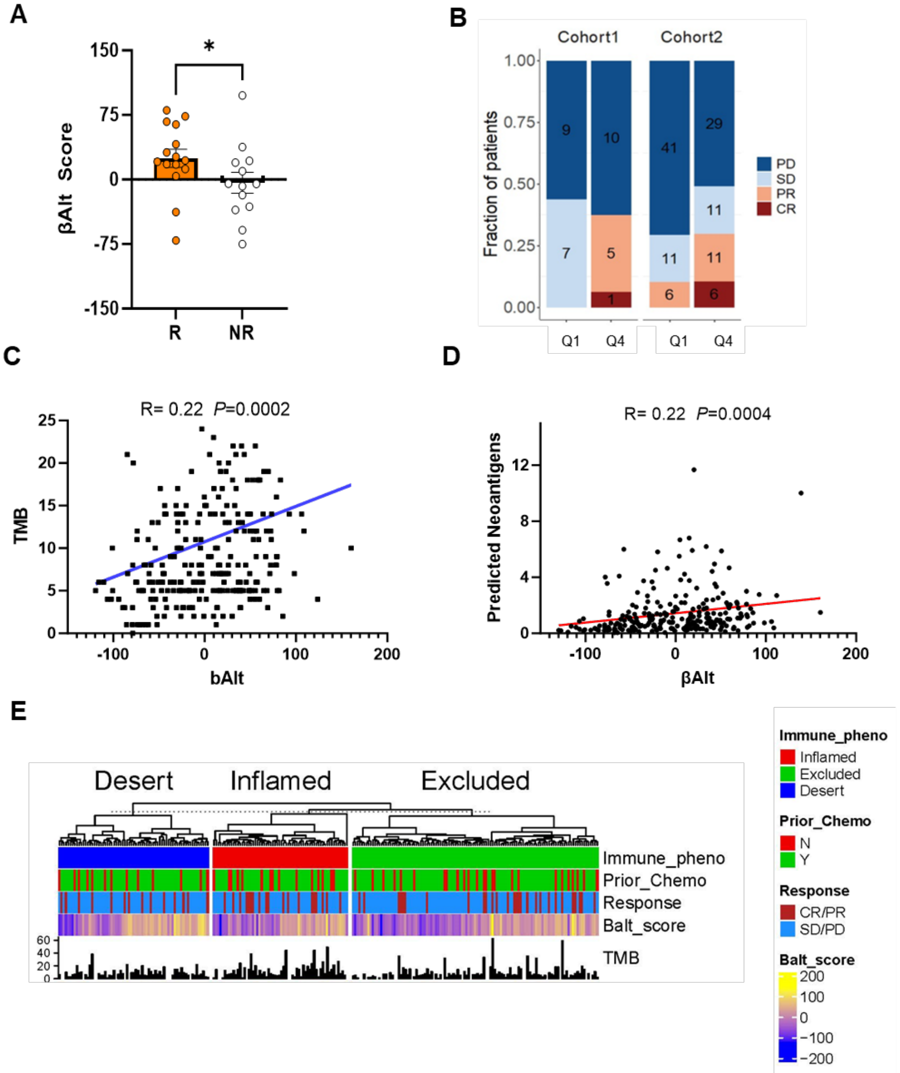

**Supplemental Fig. 2. High βAlt correlates with the immune composition in a tissue-agnostic manner. (A)** Unsupervised hierarchical clustering heatmap of type I interferon signature of IMvigor210 metastatic bladder patient above with βAlt score (red=positive, teal=negative). **(B)** Unsupervised hierarchical clustering heatmap of TeIS for IMVigor 210 as a function of assigned lymphocyte distribution [Mariathasan, 2018]. **(C)** Unsupervised hierarchical clustering heatmap of TeIS for TCGA bladder cancer (BLCA) dataset (n=405) annotated with βAlt scores above and TMB below. The immune-rich cluster is designated by a green box, the immune-poor cluster is designated by a purple box. **(D)** TeIS signature heatmap for TCGA breast cancer (BRCA) dataset (N=1082) as described above. **(E)** Expression of cytotoxic T cell markers is significantly anti-correlated with βAlt score across data sets. PCC R indicated and P values are all <0.0001.

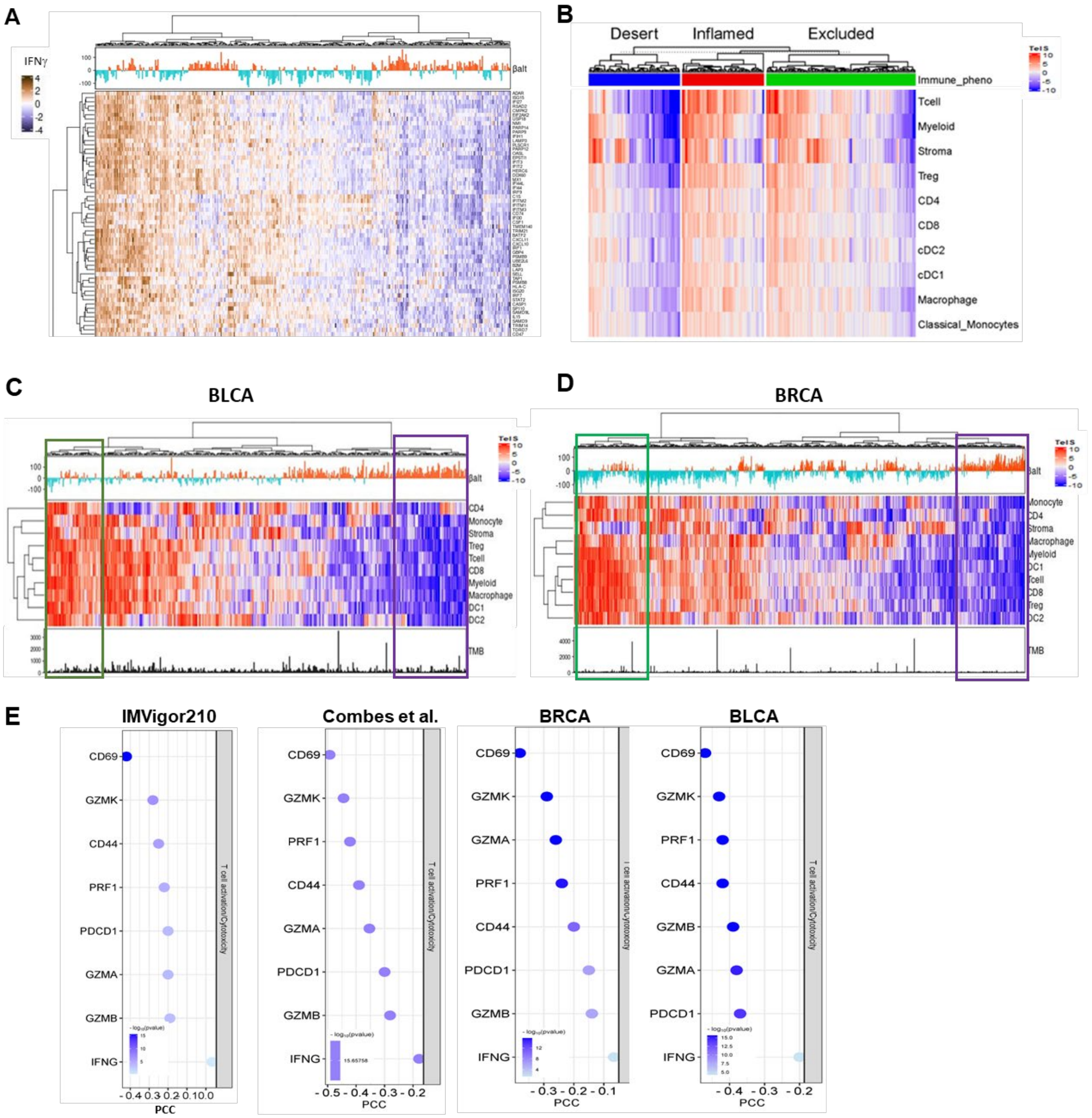

**Supplemental Fig. 3. High βAlt, immune poor tumors respond to ICB by converting to immune rich. (A)** Unsupervised hierarchical clustering heatmap of TeIS for TISMO untreated murine tumor data (n=384). **(B)** βAlt scores of TISMO tumors classified as immune-rich (n=118) and immune-poor (n=99). Wilcoxon test. **(C)** Unsupervised hierarchical clustering heatmap of TeIS signatures annotated above with βAlt scores of mTDT mammary tumors (n=70). **(D)** βAlt scores of mTDT classified as immune-rich (n=23 and immune-poor (n=13). Unpaired T test. **(E)** Tumor growth curves of mTDT compiled from different treatment arms. Based on tumor growth 7 days post treatment, mice are designated responders (R, black, n=52) and non-responders (NR, orange, n=52). **(F)** Experimental schematic of tissue collection and analysis for multispectral flow cytometry. **(G)** Distribution of tumor infiltrating CD45+ immune cell types. Percentages are derived from gating individual t-SNE islands for each treatment. **(H)** t-SNE of tumor immune cell composition from NR tumors following treatment with IgG isotype, anti-PD-L1, TGFβi, anti-PDL1+TGFβi, 10 Gy RT, RT+anti-PD-L1, RT+TGFβi and RT+TGFβi+anti-PD-L1. **(I-K)** Percentage of circulating live cells **(I)** CD8 T cells, **(J)** CD4 T cells, and **(K)** NK cells between R (n=12) and NR (n=10). Data are depicted as mean ± SEM, and unpaired t-tests. Statistical significance is denoted as follows *$P$<0.05, ** $P$<0.01

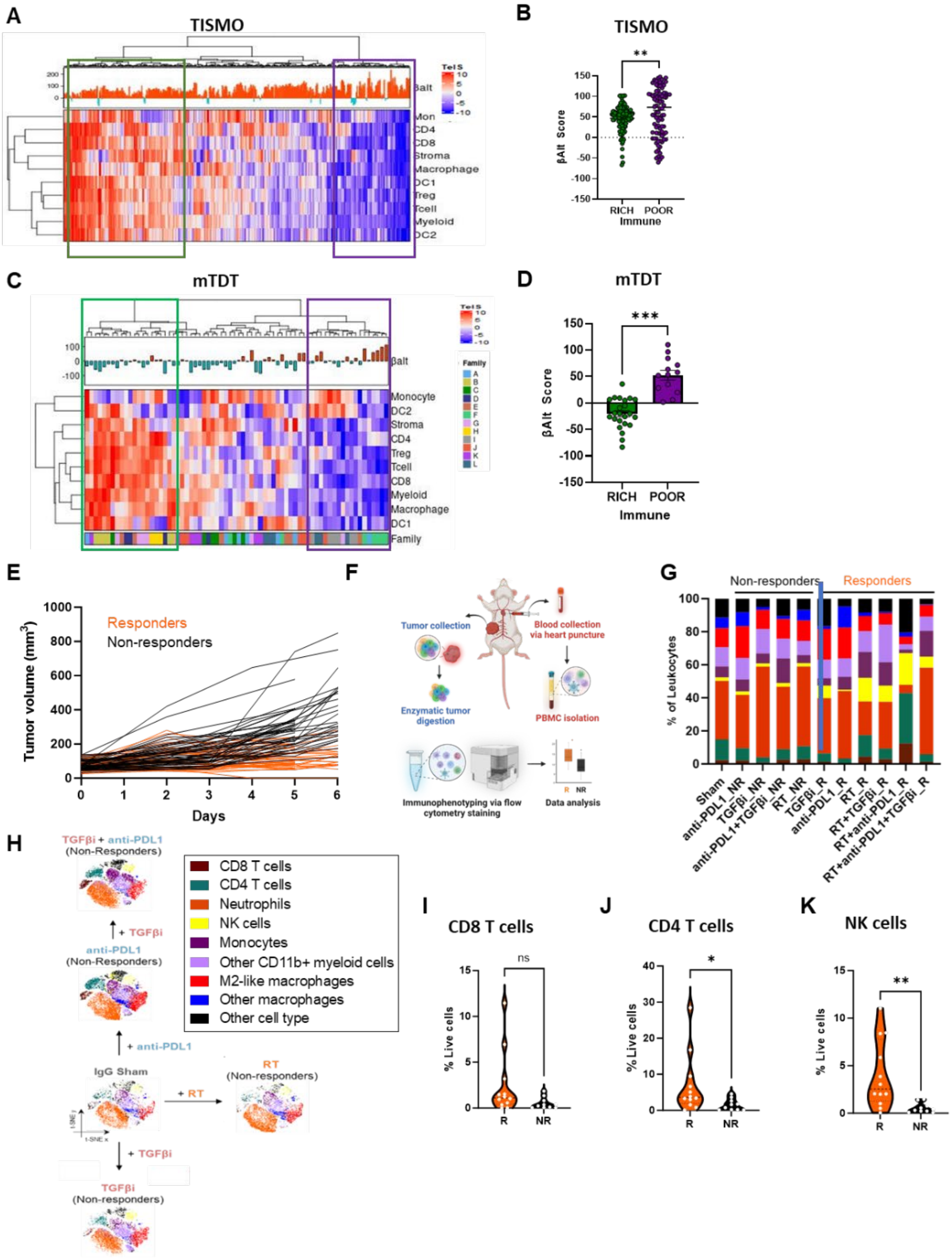

**Supplemental Fig. 4. NK cells mediate the response of βAlt high, immune-poor tumors to ICB therapy. (A)** Percent CD45+ intratumoral NK cells between mTDT R (n=13), NR (n=10) and sham (n=8). **(B)** Percent NK cell Ki67+ cells R (n=13), NR (n=10) and sham (n=8) in tumor (left). Representative fluorescence histogram (right). Percetanges of live circulating **(C) %**NK cells, **(D)** %Ki67+ NK cell cells and **(E)** %NK cell CD49b-CD40a+ ILCs of mice treated with IgG isotype without (n=2) or with NK cell depletion (n=5), RT without (n=5) or with (n=5) NK cell depletion and RT+TGFβi+anti-PDL1 without (n=5) or with (n=5) NK cell depletion. Data are mean ± SEM, each point represents a mouse, and one-way ANOVA with multiple comparisons. P-values are depicted as: * *P*<0.05, ** *P*<0.01, *** *P*<0.001. **(F)** Unsupervised hierarchical clustering heatmap of TeIS scores annotated above with βAlt scores of Riaz et al. melanoma patients treated with anti-PD1 (n=110). Bar below indicates responders (brown, n=18) and non-responders (black, n=92), and paired pretreatment biopsy (black) and on-treatment (white). **(G)** As in E but consisting only of responders with paired biopsies (n=18). Color bar below designates individuals. Note high βAlt, immune poor patients indicated by red and yellow boxes shift to immune rich. **(H)** Heatmap depicting gene expression of NK cell ligands (left) and receptors (right) high βAlt ICB responders pre (n=2) and on (n=2) treatment of melanoma patients. P values are depicted as: *** *P*<0.001, **** *P*<0.0001.

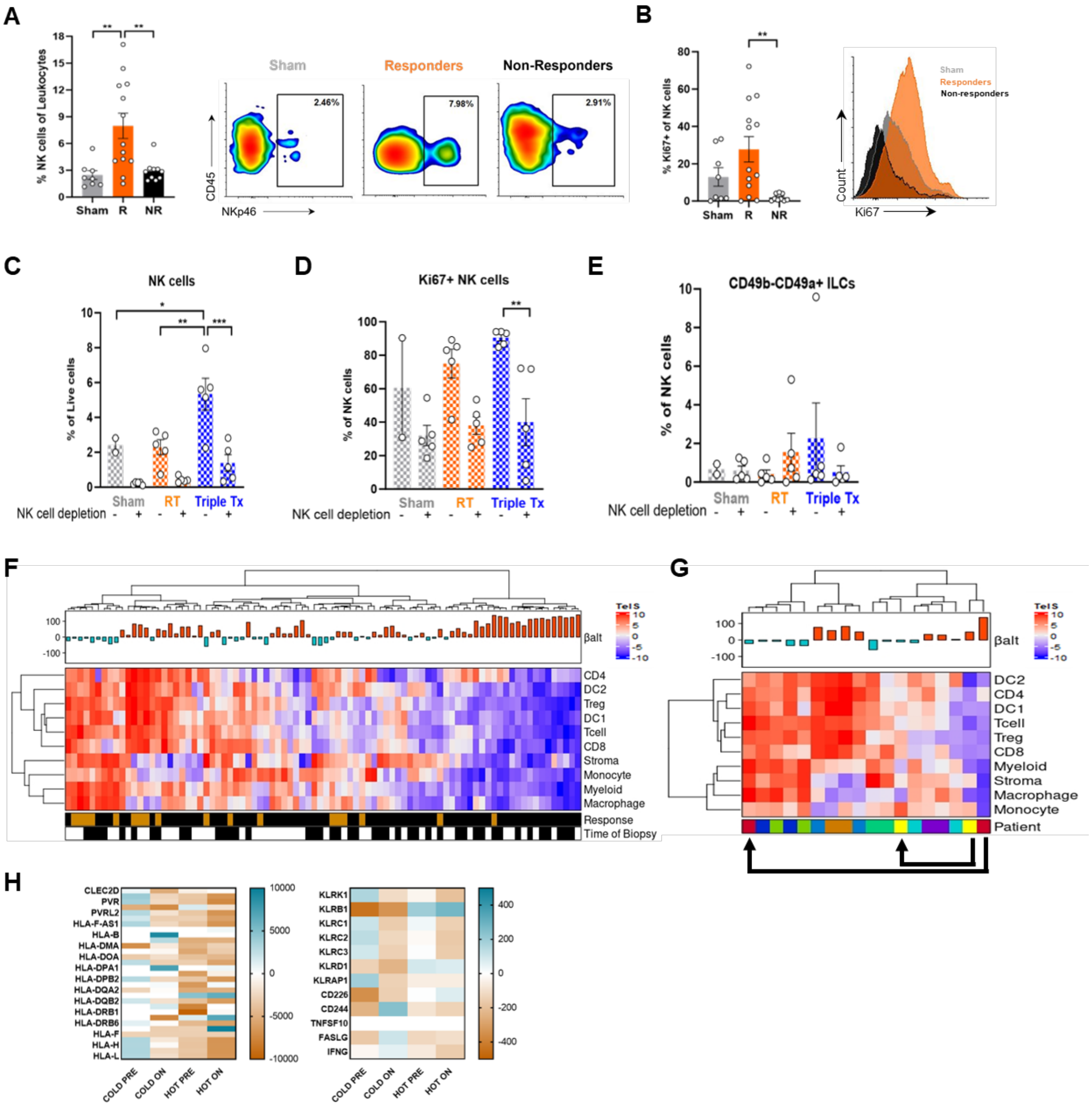

### Supplemental Table 1. Flow Cytometry Antibodies

| REAGENT or RESOURCE | SOURCE | IDENTIFIER |
|---|---|---|
| Antibodies | | |
| Rat PerCP anti-mouse CD45 monoclonal antibody | Biolegend | BioLegend Cat# 103130, RRID:AB_893339 |
| Rat Brilliant Violet 421(TM) anti-mouse CD335 (NKp46) monoclonal antibody | Biolegend | BioLegend Cat# 137612, RRID:AB_2563104 |
| Brilliant Violet 650(TM) anti-mouse/human CD11b monoclonal antibody | Biolegend | BioLegend Cat# 101259, RRID:AB_2566568 |
| Armenian hamster Spark Blue(TM) 550 anti-mouse CD11c monoclonal antibody | Biolegend | BioLegend Cat# 117366, RRID:AB_2860634 |
| Rat PerCP/Cyanine5.5 anti-mouse I-A/I-E monoclonal antibody | Biolegend | BioLegend Cat# 107625, RRID:AB_2191072 |
| Rat APC/Cyanine7 anti-mouse Ly-6C monoclonal antibody | Biolegend | BioLegend Cat# 128026, RRID:AB_10640120 |
| Rat Brilliant Violet 711(TM) anti-mouse CD317 (BST2, PDCA-1) monoclonal antibod | Biolegend | BioLegend Cat# 127039, RRID:AB_2832459 |
| Rat PE/Cyanine7 anti-mouse CD25 monoclonal antibody | Biolegend | BioLegend Cat# 102016, RRID:AB_312865 |
| Syrian hamster FITC anti-mouse Ly-49C/F/I/H monoclonal | Biolegend | BioLegend Cat# 108205, RRID:AB_2254372 |
| Rat PE anti-mouse CD215 (IL-15Ralpha) Monoclonal antibody | Biolegend | BioLegend Cat# 153503, RRID:AB_2721341 |
| Rat Alexa Fluor(R) 700 anti-mouse Ki-67 monoclonal antibody | Biolegend | BioLegend Cat# 652419, RRID:AB_2564284 |
| Rat Pacific Blue anti-mouse Perforin monoclonal antibody | Biolegend | BioLegend Cat# 154311, RRID:AB_2922481 |
| Armenian hamster Alexa Fluor(R) 647 anti-mouse CXCL9 (MIG) monoclonal antibo | Biolegend | BioLegend Cat# 515606, RRID:AB_1877135 |
| Rat Spark YG(TM) 593 anti-mouse CD14 monoclonal antibody | Biolegend | BioLegend Cat# 150107, RRID:AB_2894630 |
| Rat Brilliant Violet 570(TM) anti-mouse Ly-6G monoclonal antibody | Biolegend | BioLegend Cat# 127629, RRID:AB_10899738 |
| Rat Brilliant Violet 785(TM) anti-mouse monoclonal F4/80 antibody | Biolegend | BioLegend Cat# 123141, RRID:AB_2563667 |
| Armenian hamster FITC anti-mouse CD80 monoclonal antibody | Biolegend | BioLegend Cat# 104706, RRID:AB_313127 |
| Brilliant Violet 605(TM) anti-mouse CD274 (B7-H1, PD-L1) monoclonal antibody | Biolegend | BioLegend Cat# 124321, RRID:AB_2563635 |
| Rat PE/Fire(TM) 700 anti-mouse CD3 monoclonal antibody | Biolegend | BioLegend Cat# 100271, RRID:AB_2876394 |
| Rat PE/Fire(TM) 640 anti-mouse CD8a monoclonal antibody | Biolegend | BioLegend Cat# 100789, RRID:AB_2860590 |
| Rat APC/Fire(TM) 810 anti-mouse CD4 monoclonal antibody | Biolegend | BioLegend Cat# 100479, RRID:AB_2860583 |
| Rat IL-2 monoclonal antibody | BD Biosciences | BD Biosciences Cat# 566363, RRID:AB_2739711 |
| Rat Alexa Fluor(R) 700 anti-mouse FOXP3 monoclonal antibody | BioLegend | BioLegend Cat# 126422, RRID:AB_2750493 |
| Rat Brilliant Violet 711(TM) anti-mouse monoclonal CD8a antibody | BioLegend | BioLegend Cat# 100759, RRID:AB_2563510 |
| Rat APC anti-mouse Perforin monoclonal antibody | BioLegend | BioLegend Cat# 154404, RRID:AB_2721465 |
| Rat PE anti-mouse CD279 (PD-1) monoclonal antibody | BioLegend | BioLegend Cat# 109104, RRID:AB_313421 |
| Rat CD206 (MMR) Monoclonal Antibody (MR6F3), eFluor 450, eBioscience | eBioscience | Thermo Fisher Scientific Cat# 48-2061-82, RRID:AB_2762721 |
| Rat FITC anti-mouse GM-CSF Monoclonal antibody | BioLegend | BioLegend Cat# 505403, RRID:AB_315379 |
| Rat Brilliant Violet 421(TM) anti-mouse IFN-gamma monoclonal antibody | BioLegend | BioLegend Cat# 505829, RRID:AB_10897937 |
| Rat Brilliant Violet 711(TM) anti-mouse TNF-alpha monoclonal antibody | BioLegend | BioLegend Cat# 506349, RRID:AB_2629800 |
| Armenian hamster APC anti-mouse CD49a monoclonal antibody | BioLegend | BioLegend Cat# 142605, RRID:AB_2562252 |
| Rat PE/Cyanine7 anti-mouse CD49b (pan-NK cells) monoclonal antibody | BioLegend | BioLegend Cat# 108921, RRID:AB_2561459 |
| Mouse PE anti-mouse CD159a (NKG2AB6) monoclonal antibody | BioLegend | BioLegend Cat# 142803, RRID:AB_10959654 |
| Rat CD19 Monoclonal Antibody (eBio1D3 (1D3)), PE-Cyanine5, eBioscience | eBioscience | Thermo Fisher Scientific Cat# 15-0193-82, RRID:AB_657672 |
| Rabbit IgG phospho-SMAD2 on serine 465/467 (Cell Signaling Cat#3108, 1:200) | Cell Signaling | UniProt ID: Q15796 |
| Secondary donkey anti-rabbit IgG (Alexa Fluor 488/555) | Invitrogen | |
| Donkey anti-mouse IgG (Alexa Fluor 488/555) | Invitrogen | |
| Rat Brilliant Violet 650(TM) anti-mouse/human CD11b monoclonal antibody | BioLegend | BioLegend Cat# 101239, RRID:AB_11125575 |
| Mouse Brilliant Violet 421(TM) anti-mouse NK-1.1 monoclonal antibody | BioLegend | BioLegend Cat# 108731, RRID:AB_10895916 |
| Rat Spark NIR(TM) 685 anti-mouse CD19 monoclonal antibody | BioLegend | BioLegend Cat# 115567, RRID:AB_2819828 |
| Armenian hamster Pacific Blue(TM) anti-mouse CD11c monoclonal antibody | BioLegend | BioLegend Cat# 117321, RRID:AB_755987 |
| Rat PE/Dazzle(TM) 594 anti-mouse CD62L monoclonal antibody | BioLegend | BioLegend Cat# 104447, RRID:AB_2566162 |
| Rat Brilliant Violet 605(TM) anti-mouse CD115 (CSF-1R) monoclonal antibody | BioLegend | BioLegend Cat# 135517, RRID:AB_2562760 |
| Rat CD4 Monoclonal Antibody (RM4-5), Brilliant Ultra Violet 805, eBioscience | Thermo Fisher Scientific | Thermo Fisher Scientific Cat# 368-0042-82, RRID:AB_2896072 |
| Rat APC anti-mouse CD274 (B7-H1, PD-L1) monoclonal antibody | BioLegend | BioLegend Cat# 124311, RRID:AB_10612935 |
| Rat Alexa Fluor(R) 488 anti-mouse Ki-67 monoclonal antibody | BioLegend | BioLegend Cat# 652417, RRID:AB_2564236 |

#### Supplemental Table 2. Flow Cytometry Gating

| Immune population | Marker combination for identification |
|---|---|
| Leukocytes | Zombie Aqua-, CD45+ |
| Total T cells | Zombie Aqua-, CD45+, CD11b-, NKp46-, CD3+ |
| CD8 T cells | Zombie Aqua-, CD45+, CD11b-, NKp46-, CD3+, CD4-, CD8+ |
| Proliferating CD8 T cells | Zombie Aqua-, CD45+, CD11b-, NKp46-, CD3+, CD4-, CD8+, Ki67+ |
| IL-2 CD8 T cells | Zombie Aqua-, CD45+, CD11b-, NKp46-, CD3+, CD4-, CD8+, IL-2+ |
| CD4 T cells | Zombie Aqua-, CD45+, CD11b-, NKp46-, CD3+, CD8-, CD4+ |
| Proliferating CD4 T cells | Zombie Aqua-, CD45+, CD11b-, NKp46-, CD3+, CD8-, CD4+, Ki67+ |
| Natural Killer (NK) cells | Zombie Aqua-, CD45+, CD3-, NKp46+ |
| Proliferating NK cells | Zombie Aqua-, CD45+, CD3-, NKp46+, Ki67+ |
| Circulating Innate Lymphoid Cells (ILC) | Zombie Aqua- blood cells, CD45+, CD3-, NKp46+, CD49b-, CD49a+ |
| Dendritic cells | Zombie Aqua-, CD45+, CD3-, NKp46-, CD11b+, Ly6G-, F4/80-, MHCII+ |
| Classical Dendritic cells type-I (cDC-1) | Zombie Aqua-, CD45+, CD3-, NKp46-, CD11b+, Ly6G-, F4/80-, MHCII+, CD11c+ |
| CXCL9 cDC-1 | Zombie Aqua-, CD45+, CD3-, NKp46-, CD11b+, Ly6G-, F4/80-, CD11c+, MHCII+, CXCL9+ |
| IL-2 cDC-1 | Zombie Aqua-, CD45+, CD3-, NKp46-, CD11b+, Ly6G-, F4/80-, CD11c+, MHCII+, IL-2+ |
| TNFa cDC-1 | Zombie Aqua-, CD45+, CD3-, NKp46-, CD11b+, Ly6G-, F4/80-, CD11c+, MHCII+, TNFa+ |
| Total PBMC | Zombie Aqua- blood cells |
| IL-2 Total PBMC | Zombie Aqua- blood cells, IL-2+ |
| TNFa Total PBMC | Zombie Aqua- blood cells, TNFa+ |
| CXCL9 Total PBMC | Zombie Aqua- blood cells, CXCL9+ |
| Neutrophils | Zombie Aqua-, CD45+, CD3-, NKp46-, CD11b+, Ly6G+ |
| B cells | Zombie Aqua-, CD45+, CD3-, NKp46-, CD19+ |
| Macrophages | Zombie Aqua-, CD45+, CD3-, NKp46-, CD11b+, Ly6C+, Ly6G-, F4/80+ |
| M2-like Macrophages | Zombie Aqua-, CD45+, CD3-, NKp46-, CD11b+, Ly6C+, Ly6G-, F4/80+, CD206+ |
| Other Macrophages | Zombie Aqua-, CD45+, CD3-, NKp46-, CD11b+, Ly6C+, Ly6G-, F4/80+, CD206- |
| Monocytes | Zombie Aqua-, CD45+, CD3-, NKp46-, CD11b+, Ly6C+ |
| Other CD11b+ myeloid cells | Zombie Aqua-, CD45+, CD3-, NKp46-, CD11b+, Ly6C-, Ly6G-, F4/80-, MHCII- |

#### Supplemental Table 3. βAlt signature representing TGFβ and alt-EJ gene lists

| TGFβ genes | alt-EJ genes |
|---|---|
| ABCG1 | APE2 |
| AMIGO2 | APEX1 |
| CA12 | ASF1A |
| CCDC99 | CDKN2D |
| CCL20 | CIB1 |
| CHRNA9 | DNA2 |
| COL4A2 | FAAP24 |
| CTGF | FANCM |
| DLC1 | GEN1 |
| DNAJB9 | HRAS1 |
| DSC2 | LIG1 |
| ENC1 | LIG3 |
| F3 | MEN1 |
| FAP | MRE11A |
| FGF2 | MSH3 |
| FN1 | MSH6 |
| HEY1 | MTH1 |
| HMGA2 | MTOR |
| ID1 | NABP2 |
| IGF2BP3 | NTHL1 |
| IGFBP3 | PALB2 |
| JAG1 | PARP1 |
| KLF4 | PARP3 |
| LAMB3 | POLA1 |
| LAMC2 | POLM |
| LARP6 | POLQ |
| LIPG | PRP19 |
| MAFF | RAD51D |
| MMD | RBBP8 |
| PDGFC | RRM2 |
| PLEK2 | RUVBL2 |
| LEXNA2 | SOD1 |
| PSTN | TIP60 |
| PSCD1 | UNG |
| RICS | WRN |
| RNF24 | XRCC1 |
| RUNX1 | |
| SAMSN1 | |
| LAMC2 | |
| SERPINE1 | |
| SERPINE2 | |
| SH2D2A | |
| SH2D4A | |
| SLC20A1 | |
| SLC22A4 | |
| TGIF1 | |
| THBS1 | |
| TMEPAI | |
| TNC | |
| TNFRAF12A | |
| VACN | |

#### Supplemental Table 4. Tumor-educated immune signatures (TeIS) gene lists

| Tcell | Myeloid | Stroma | CD4 | CD8 | Treg | Macrophage | Classical Monocytes | cDC1 | cDC2 |
|---|---|---|---|---|---|---|---|---|---|
| CD40LG | CLEC10A | CDH11 | BACH2 | FASLG | FOXP3 | APOE | S100A8 | IDO1 | FCER1A |
| TBX21 | CD1E | DCN | TRABD2A | CD8A | IL2RA | TREM2 | SL00A9 | C1ORF54 | CD1C |
| SH2D1A | CD1C | PDGFRA | IL7R | SETBP1 | CD80 | C1QB | VCAN | XCR1 | CD1E |
| PYHIN1 | VSIG4 | COL1A2 | HDAC4 | CTSW | CD177 | C1QC | FCN1 | CLEC9A | CD1D |
| ZNF831 | CD33 | ISLR | NR3C2 | APOBEC3( | LAIR2 | VSIG4 | LYZ | BATF3 | CLEC10A |
| CD6 | CD300LB | COL1A1 | ADD3 | APOBEC3( | CCR8 | | | | |
| THEMIS | MS4A7 | FNDC1 | PABPC1 | BTNL8 | CCL22 | | | | |
| UBASH3A | LY86 | BGN | PABPC3 | HLA-DPB1 | TNFRSF13B | | | | |
| TRAT1 | CLEC5A | COL5A2 | MFHAS1 | ULBP3 | TNFRSF18 | | | | |
| EOMES | LILRB4 | POSTN | DSC1 | RAD51 | | | | | |
| GRAP2 | FCER1A | PCDH18 | SELL | WNT9A | | | | | |
| ZAP70 | LIL4B2 | ADAMTS1 | SESN1 | HLA-DMA | | | | | |
| SIRPG | CSF1R | MXRA5 | | CERS5 | | | | | |
| ICOS | LILRA1 | SULF1 | | CD8B | | | | | |
| FASLG | ADORA3 | EDNRA | | NKG7 | | | | | |
| CD8A | MPEG1 | PRRX1 | | FAM156A | | | | | |
| CD8B | FCN1 | COL3A1 | | SNF696 | | | | | |
| ITK | RNASE6 | THY1 | | MCM5 | | | | | |
| GZMA | FPR3 | LUM | | DPF3 | | | | | |
| KLRK1 | CYBB | COL12A1 | | TTC24 | | | | | |
| GZMH | MS4A4A | | | YARS | | | | | |
| GZMK | MS4A4E | | | EBP | | | | | |
| CD3D | OLR1 | | | TRPS1 | | | | | |
| CD3E | CD163 | | | GZMH | | | | | |
| CD3G | WDFY4 | | | VCAM1 | | | | | |
| | SIGLEC7 | | | LAG3 | | | | | |
| | MS4A14 | | | CRIM1 | | | | | |
| | SIGLEC1 | | | NAA40 | | | | | |
| | CD300E | | | GRIK4 | | | | | |
| | | | | PSMB9 | | | | | |
| | | | | HLA-DMB | | | | | |
| | | | | SERP2 | | | | | |
| | | | | HOXB4 | | | | | |
| | | | | CASP7 | | | | | |
| | | | | TMCC2 | | | | | |
| | | | | ARPC5L | | | | | |
| | | | | MCTP2 | | | | | |
| | | | | LYST | | | | | |
